# In Luminal A breast cancer, bulk *FOXC1* reports the tumour vasculature, while a proliferation-independent basal-lineage axis marks the tumour cells

**DOI:** 10.64898/2026.08.11.744143

**Authors:** Daniel E. Yehoshua, Joseph C. Bingham

## Abstract

*FOXC1* is widely interpreted as a tumour-cell driver of basal identity and immune phenotype in breast cancer, including within the Luminal A (LumA) subtype, where its bulk expression tracks basal-like features and immune infiltration. Because LumA tumours are compositionally heterogeneous, whether this reflects malignant-cell biology or the surrounding microenvironment has not been resolved, and the distinction changes how the biomarker should be read. Combining purity- and compartment-adjusted partial correlations across three LumA cohorts on two platforms (TCGA, *n* = 571; METABRIC, *n* = 700; SCAN-B, *n* = 1,540) with single-cell data from an ER-positive atlas (GSE176078; 100,064 cells), we show that bulk *FOXC1* is principally a readout of the tumour vasculature: roughly 90% of *FOXC1* transcripts arise from endothelial and perivascular cells and 1.8% from malignant epithelium, and *FOXC1* is detected in 28% of endothelial versus 0.5% of malignant cells. Its coupling to adaptive-immune and tertiary-lymphoid-structure programmes collapses under vascular adjustment, whereas basal cytokeratins (*KRT5*, *KRT14*, *KRT17*) and *TP63* retain a residual, population-level basal-lineage signal, the Centaur axis, that is orthogonal to proliferation and to established risk tools; an apparent survival association is explained by age. These findings reassign a presumed tumour-cell programme to the vessel wall, recover a genuine basal-lineage axis within LumA tumour cells, and show that compartment-aware analysis is needed to interpret bulk transcriptomic biomarkers in compositionally heterogeneous tumours.

**Impact statement:** In Luminal A breast cancer, single-cell and compartment-adjusted analysis shows that bulk *FOXC1* reports the tumour vasculature rather than the malignant cells, while a genuine basal-lineage axis persists in the tumour cells.

## Introduction

### Clinical gap in Luminal A

The Luminal A subtype represents approximately 40–50% of breast cancer diagnoses [1, 2, 3, 4], and although generally favourable prognostically, approximately 15–20% of patients experience disease relapse despite appropriate endocrine therapy [5, 6]. Luminal A tumours are characterised as immunologically cold, exhibiting minimal T-cell infiltration, low tumour mutational burden, and rare tertiary lymphoid structure (TLS) formation [7, 8]; consequently, they have been largely excluded from immune-checkpoint trials [9, 10], and current ICB biomarkers stratify poorly within the ER-positive population [11, 12, 26]. An unmet need persists for mechanism-based molecular stratification within LumA.

A recurring difficulty is that candidate LumA-relevant transcription factors identified from bulk RNA analyses – including *FOXC1* – have well-established activity in vascular or stromal compartments [14, 13]. Bulk-level correlations between such genes and immune programmes therefore have two admissible explanations: a genuine tumour-cell-intrinsic effect, or co-variation with the size of the vascular and cancer-associated fibroblast (CAF) compartments in which the same gene is also expressed. These are not the same claim and have different clinical implications, yet the distinction has not been systematically enforced in prior work on *FOXC1* in Luminal A. We adopt purity- and stroma-adjusted partial correlations as the minimum analytical standard and re-characterise the *FOXC1* programme in Luminal A from first principles.

### FOXC1 and the Centaur concept

*FOXC1* is a forkhead-box transcription factor with roles in vascular, ocular, and mesenchymal development [13, 14], and in the adult mammary gland is expressed in *CD44*^+^ progenitor cells [42] and silenced during luminal differentiation via *EZH2*-mediated H3K27me3 [15, 16]. It is a PAM50 basal-defining gene [1]; elevated *FOXC1* in basal-like disease correlates with poor outcomes [17, 18], while in Luminal B it is paradoxically anti-metastatic via a distinct *EZH2*-mediated mechanism [15], indicating that *FOXC1*’s effect is fundamentally context-dependent. In cell-line models, ectopic *FOXC1* overexpression represses ER*α* at levels characteristic of basal-like disease [17]; whether comparable, if quantitatively smaller, effects occur at the modest *FOXC1* elevations observed within the Luminal A expression range has not been examined directly.

We term the axis along which the basal-lineage programme is activated across Luminal A tumours the *molecular Centaur*, evoking the mythological creature whose human and equine anatomy coexist: a single entity in which two lineage programmes are jointly present. As we show, this coexistence is a property of the tumour *population* – some Luminal A tumours engage the basal-lineage programme more than others – rather than of individual dual-lineage cells. Concretely, the Centaur state is a graded, population-level co-elevation of a tumour-cell basal-lineage programme on a retained luminal-identity background within PAM50-classified Luminal A tumours. It is distinct from epithelial–mesenchymal transition (a mesenchymal, not basal-epithelial, programme), from admixture of separate luminal and basal-like populations (the signal is graded, not a two-population mixture), from classical basal-like disease (Centaur tumours remain Luminal A by PAM50, far below basal-range *FOXC1*), and from within-cell lineage plasticity (unsupported by the single-cell data here). We characterise the axis with the following aims: (i) establish its molecular signature and confirm it occupies a graded within-Luminal A rather than boundary-classification position; (ii) apply purity- and stroma-adjusted partial correlations to determine which associated microenvironmental programmes reflect tumour-compartment biology versus stromal-vascular co-variation; (iii) benchmark *FOXC1* against alternative basal-lineage anchors under identical compartment adjustment; (iv) determine the axis’s relationship to existing proliferation-based clinical tests and immune signatures and evaluate its prognostic independence; and (v) resolve the cellular source of the bulk *FOXC1* signal at single-cell resolution and test whether the basal-lineage axis reflects within-cell co-expression.

## Results

### Basal-lineage activation defines a graded axis within Luminal A

If a hybrid lineage state exists within Luminal A, the transcription factor that drives it should vary across tumours without splitting them into discrete groups – a continuous axis of basal-programme activation rather than a hidden subtype. This is what we observed. Across 571 PAM50-assigned Luminal A tumours from TCGA-BRCA (cohort characteristics in Table 1), *FOXC1* expression formed a single continuous, approximately Gaussian distribution with no evidence of a discrete high-expressing subpopulation (Hartigan’s dip test *p* = 0.97; a one-component model was favoured over a two-component mixture). *FOXC1* levels in Luminal A spanned a wide range but remained well below those of basal-like tumours: only about 1% of Luminal A tumours reached even the bottom decile of Basal *FOXC1* expression. The lineage state we describe therefore arises from modest *FOXC1* elevations within the luminal range, not from tumours that have crossed into basal-level expression. The divergently transcribed lncRNA *FOXCUT* tracked *FOXC1* closely (*ρ* = 0.68), consistent with the established regulatory relationship in which *FOXCUT* sustains *FOXC1* expression [19, 20, 21, 22, 23].

**Table 1:** Cohort characteristics stratified by Centaur status. Centaur/non-Centaur groups defined by a median split of the basal-epithelial lineage axis within Luminal A in each cohort (Methods; the *FOXC1* single-gene values are shown for reference). Age and follow-up shown as median (IQR). Stage distribution shown as percentage of Luminal A stratum. Radiation and hormone therapy shown as percentage of subgroup. TCGA values extracted from TCGA-BRCA Pan-Cancer Atlas via cBioPortal (n=499 LumA; see note below); METABRIC values from full cBioPortal download. TCGA OS events shown per Centaur group (annotated n=499 subset); the paper’s primary radiation-naive survival stratum had 21 events in 218 patients.

| Characteristic | TCGA-BRCA LumA |  | METABRIC LumA |  |
| --- | --- | --- | --- | --- |
|  | Non-Centaur<br>(n = 250) | Centaur (n = 249) | Non-Centaur<br>(n = 350) | Centaur (n = 350) |
| Age at diagnosis, yr | 62.0 (51–71) | 56.0 (47–65) | 64.0 (53.8–72.5) | 62.6 (53.0–71.6) |
| Stage (%) |  |  |  |  |
| I | 22.0 | 22.8 | 30.3 | 33.1 |
| II | 58.5 | 51.6 | 42.0 | 39.7 |
| III | 19.5 | 25.6<sup>†</sup> | 2.3 | 5.1<sup>†</sup> |
| Unstaged/missing | 0.0 | 0.0 | 25.4 | 22.1 |
| OS events, n (%) | 31 (12.4%) | 26 (10.4%) | 151 (43.1%) | 143 (40.9%) |
| DSS events, n | 14 | 13 | 131 total (LumA) |  |
| Median follow-up, mo | 23.9 | 33.9 | 140 | 127 |
| Treatment (%) |  |  |  |  |
| Radiation (yes) | 51.8 | 57.9 | 57.1 | 50.9 |
| Hormone therapy (yes) | — | — | 67.7 | 68.6 |
| Chemotherapy (yes) | — | — | 5.1 | 10.9<sup>†</sup> |
| *FOXC1*, median log<sub>2</sub> | 6.30 | 7.73 | 7.53 | 8.43 |
| *FOXC1* platform IQR | 6.3–7.7 (RSEM log<sub>2</sub>) |  | 7.53–8.43 (microarray) |  |

To operationalise the Centaur phenotype as a lineage programme rather than a single gene, and to guard against any single high-variance keratin dominating, we defined a basal-epithelial lineage score as the mean per-gene z-score across a panel of tumour-cell-intrinsic basal markers (*KRT5*, *KRT14*, *KRT17*, *KRT6B*, *EGFR*, *TP63*, *CDH3*, *MIA*, *SFRP1*, *S100A2*, and *FOXC1*). We deliberately excluded myoepithelial, EMT, and stromal genes (e.g. *ACTA2*, *MYH11*, *VIM*, *NGFR*) from this panel: including them would make the score partly a readout of stromal content and would mechanically inflate the downstream stromal and immune associations we report below. Within Luminal A, *FOXC1* correlated with the remaining basal programme even when *FOXC1* itself was excluded from the score (*ρ* = 0.46), confirming that *FOXC1* elevation marks activation of a coordinated basal programme rather than isolated single-gene expression. This basal-epithelial axis, which defines Centaur-high versus Centaur-low tumours throughout, correlated strongly with *FOXC1* alone (*ρ* = 0.75) but is not identical to it, and provides a lineage-anchored definition that is not confounded with the immune or stromal compartments. Hereafter, “Centaur phenotype” refers to the tumour-level axis defined by this basal-epithelial lineage composite score, and the composite score is its operational measure.

### FOXC1’s bulk immune coupling in LumA is an endothelial-mediated signal, not a tumour-cell programme

We first asked what accompanies *FOXC1* elevation in the Luminal A tumour microenvironment. In unadjusted analyses, *FOXC1* correlated with a coordinated adaptive-immune programme spanning the homeostatic chemokines *CCL19* and *CCL21* [25], B-cell markers, cytotoxic effectors, and germinal-centre follicular dendritic cell markers (Supplementary Table S1), a pattern that would ordinarily be described as an organised, TLS-linked adaptive-immune signature. The correlations replicated in direction in METABRIC (19/37 immune genes at FDR *<* 0.05), though at attenuated magnitudes.

*FOXC1*, however, has canonical vascular activity, and its bulk-level correlations must therefore be evaluated against the possibility that they reflect co-variation with the vascular and CAF compartments rather than tumour-cell biology. We tested this directly using partial-Spearman correlations of *FOXC1* with each immune gene, adjusted for a joint composite of vascular (*CDH5*, *PECAM1*, *VWF*, *ACKR1*, *SELP*, *SELE*, *PLVAP*, *VCAM1*, *ICAM1*) and CAF (*PDGFRA*, *PDGFRB*, *ACTA2*) markers computed within the Luminal A stratum. Using these predefined vascular+CAF composites, adjustment substantially attenuated – and in most categories reversed – the *FOXC1*-immune correlations (Fig. 2). In TCGA, the median unadjusted *FOXC1* correlation across TLS-organising chemokines dropped from *ρ* = +0.41 to *ρ* = −0.00 under joint vascular+CAF adjustment; B-cell markers from +0.37 to +0.04; cytotoxic effectors from +0.38 to −0.03; and mature dendritic-cell markers from +0.38 to +0.01. METABRIC showed the same collapse under adjustment, with attenuated magnitudes and residuals landing at or just below zero (TLS +0.26 → −0.04; B-cell +0.14 → −0.02; cytotoxic +0.18 → −0.11; mature DC +0.19 → −0.04). No immune category retained any FDR-significant positive *FOXC1* correlations under the adjusted model in either cohort (Fig. 2; Supplementary Table S3).

**Figure 1.**
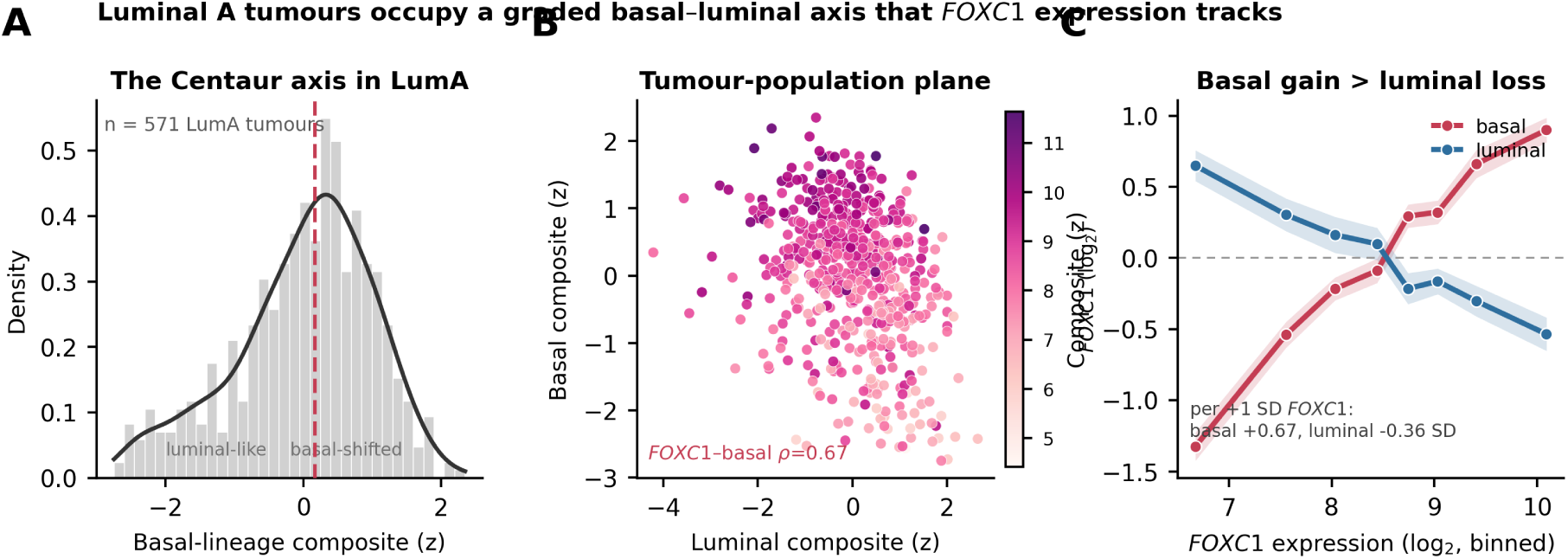
The Centaur axis is a graded basal–luminal continuum within Luminal A that *FOXC1* expression tracks (TCGA, *n* = 571). (A) Distribution of the basal-lineage composite (the Centaur axis) across Luminal A tumours: a continuous, unimodal gradient (dashed line, median) from luminal-like to basal-shifted, with no discrete subpopulation. (B) Luminal-identity versus basal-lineage composite plane, each point a tumour coloured by *FOXC1* expression; Luminal A tumours occupy a continuous plane and *FOXC1* tracks the basal axis (*ρ* = 0.67). (C) Across the *FOXC1* expression range the basal-lineage composite rises while the luminal-identity composite declines more modestly (per +1 SD *FOXC1*: basal +0.67 SD, luminal −0.36 SD), quantifying basal-programme gain in excess of luminal-identity attenuation.

**Figure 2.**
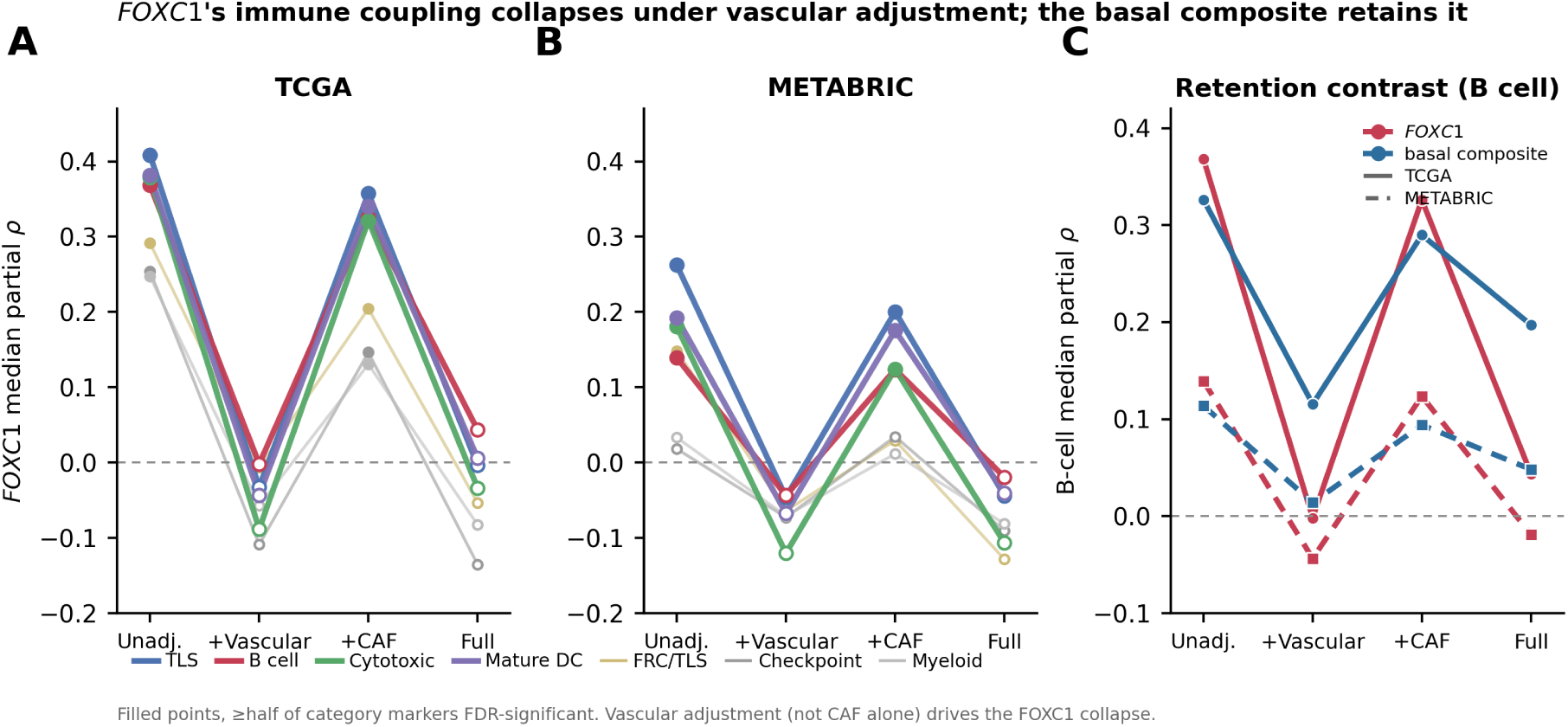
Compartment adjustment recasts *FOXC1*’s immune coupling as a vascular readout, while the basal-lineage composite retains it. Category-level median partial Spearman correlations across four nested compartment-adjustment models within Luminal A (Unadjusted; + Vascular composite; + CAF composite; joint Vascular+CAF, “Full”), lines connecting the median trajectory per immune category. (A,B) Single-gene *FOXC1* as the axis in TCGA (*n* = 571) and METABRIC (*n* = 700): the unadjusted couplings, the organised adaptive-immune and stromal programmes that replicate across both platforms (Supplementary Table S1), attenuate to near-zero or reverse under vascular and Full adjustment, with vascular adjustment (not CAF alone) driving the collapse. (C) B-cell median partial correlation for *FOXC1* versus the basal-lineage composite across the same models in both cohorts: the basal composite retains substantial residual coupling (50–60% of unadjusted magnitude in TCGA) where *FOXC1* collapses. Filled points, ≥half of category markers FDR-significant. Per-gene resolution in Supplementary Fig. S2; per-category values in Supplementary Table S3.

The magnitude of this attenuation, and its consistency across cohorts and immune categories, argues that the bulk-level *FOXC1*-immune signal in Luminal A is statistically absorbed by vascular and CAF composition; for *FOXC1* specifically, the single-cell attribution to the vessel wall (below) confirms this reflects compartment co-variation rather than mediation of a tumour-cell effect. Consistent with this, the strongest *FOXC1* correlations anywhere in the dataset are endothelial rather than immune (e.g. *CDH5 ρ* = 0.58, *ACKR1 ρ* = 0.53, *VWF ρ* = 0.57, *PECAM1 ρ* = 0.43 in METABRIC; Supplementary Table S1), exceeding every immune-gene correlation and matching *FOXC1*’s known role in arterial and venular specification.

The attenuation is not uniform over the composite, and its structure is informative. Refitting the *FOXC1*–immune partial correlations under alternative vascular specifications (Supplementary Table S9; Fig. 4A) shows the collapse is carried specifically by the leukocyte-adhesion/trafficking markers (*ACKR1*, *VCAM1*, *ICAM1*, *SELP*, *SELE*): under a structural-only endothelial composite (*CDH5*, *PECAM1*, *VWF*, *PLVAP*) that omits them, *FOXC1*’s B-cell coupling returns (median partial *ρ* = +0.19, 6/6 FDR-significant in TCGA; +0.08, 4/6 in SCAN-B), and adding a pericyte composite does not remove it. Because *FOXC1* is ∼90% endothelial (below) and those adhesion molecules mark the immune-recruiting, adhesion-competent endothelium *FOXC1* itself labels, this is the expected signature of an endothelial-*mediated* association rather than a spurious one: adjusting for the adhesion machinery removes *FOXC1*’s immune coupling because it removes the vascular phenotype through which that coupling runs. Adding the single-cell-derived myoepithelial score drives *FOXC1*’s residual below zero (B-cell *ρ* = +0.04 → −0.05 in TCGA, −0.00 → −0.07 in SCAN-B), leaving no epithelial remainder (Fig. 4B). *FOXC1*’s bulk immune signal is thus fully partitioned into non-malignant compartments – adhesion-competent vasculature and myoepithelium – consistent with *FOXC1* marking the immune-recruiting vessel wall rather than exerting a tumour-cell effect.

### Single-cell compartment attribution confirms FOXC1’s vascular cellular source in Luminal A

The partial-correlation evidence above is indirect: it shows the bulk *FOXC1*–immune signal is statistically explained by vascular and CAF composition, but bulk data cannot assign the *FOXC1* transcript itself to a cell type. We resolved this directly in the Wu et al. single-cell atlas (GSE176078; 100,064 cells from 26 tumours) [34]. In ER^+^ tumours, malignant epithelial cells contributed only 1.8% of all *FOXC1* transcripts, versus 73.8% from endothelial cells and 16.2% from perivascular cells – 90.0% from the vessel wall – with cancer-associated fibroblasts contributing a further 6.0%. Across the annotated compartments *FOXC1*’s distribution was near-identical to the endothelial markers *PECAM1*, *CDH5* and *VWF* (Pearson *r* = 0.98) and anticorrelated with the epithelial marker *EPCAM* (*r* = −0.20); this reflected genuine expression rather than dropout, *FOXC1* being detected in 28.0% of ER^+^ endothelial cells (per-patient tumour/endothelial log_2_ ratio −7.28). The attribution was Luminal A-specific: restricting to the five lowest-proliferation (LumA-like) ER^+^ tumours reproduced it (endothelial share 77.7%; malignant share 4.3%). It was also graded across subtype – the malignant contribution to total *FOXC1* was 1.8% in ER^+^ and 2.6% in HER2^+^ but 43.3% in TNBC – consistent with the established expression of *FOXC1* in basal-like cancer cells and confining a tumour-intrinsic *FOXC1* signal to basal disease. Single-cell resolution therefore establishes directly what the compartment-adjusted bulk analysis inferred: in Luminal A, *FOXC1* is expressed by the vessel wall, not the malignant epithelium (Fig. 3A–C). This attribution is robust to ambient RNA: DecontX decontamination (median per-cell contamination 1.6%) left the malignant *FOXC1* share essentially unchanged (2.0% → 1.8%) and the vessel-wall share marginally higher (91.9% → 92.9%), indicating that ambient contamination, if anything, mildly inflated rather than created the small malignant fraction (Methods).

**Figure 3.**
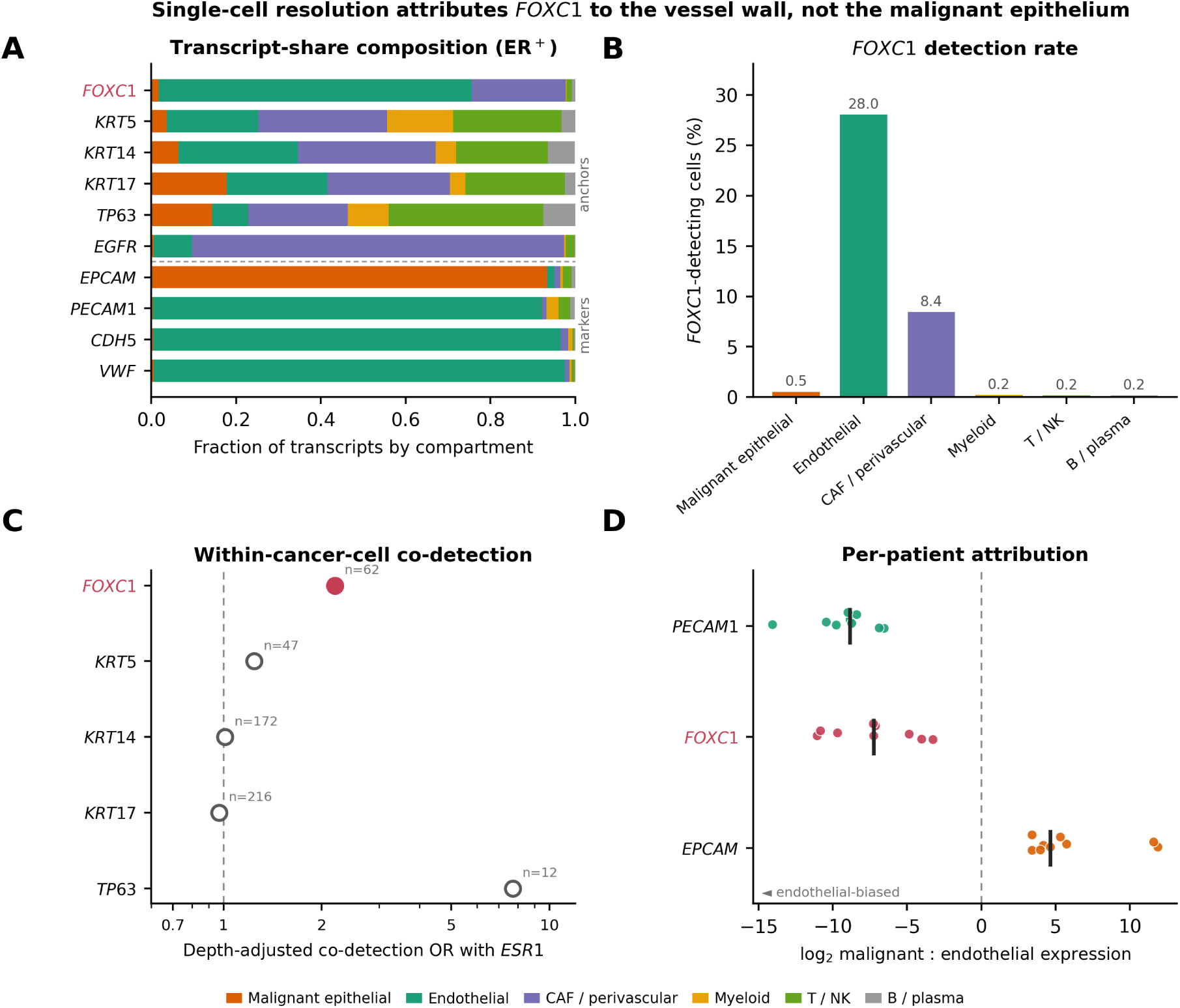
Single-cell compartment attribution places *FOXC1* in the vessel wall, not the malignant epithelium (GSE176078, ER^+^). (A) Compartment composition of each anchor’s transcripts; *FOXC1* groups with the endothelial reference markers *PECAM1*/*CDH5*/*VWF* and away from epithelial *EPCAM*, with ∼1.8% of *FOXC1* transcripts from malignant cells and ∼90% from the vessel wall. (B) *FOXC1* detection rate by compartment: 28% of endothelial cells versus 0.5% of malignant cells. (C) Depth-adjusted odds ratios for within-cancer-cell co-detection of each basal anchor with *ESR1*; keratin/*TP63* anchors straddle OR=1 (n.s.: *KRT5* 1.24, *KRT14* 1.01, *KRT17* 0.97, *TP63* 7.73 on 12 cells), and *FOXC1*’s positive OR (2.20) persists under DecontX ambient correction (2.36) so is not ambient spillover but rests on the ∼0.5% of malignant cells detecting *FOXC1* and is underpowered; *FOXC1* is the reclassified vascular gene, not a basal anchor. Filled points, *p <* 0.05. (D) Per-patient log_2_ malignant:endothelial expression ratio; *FOXC1* tracks the endothelial reference *PECAM1* (endothelial-biased in every patient), opposite to epithelial *EPCAM*.

**Figure 4.**
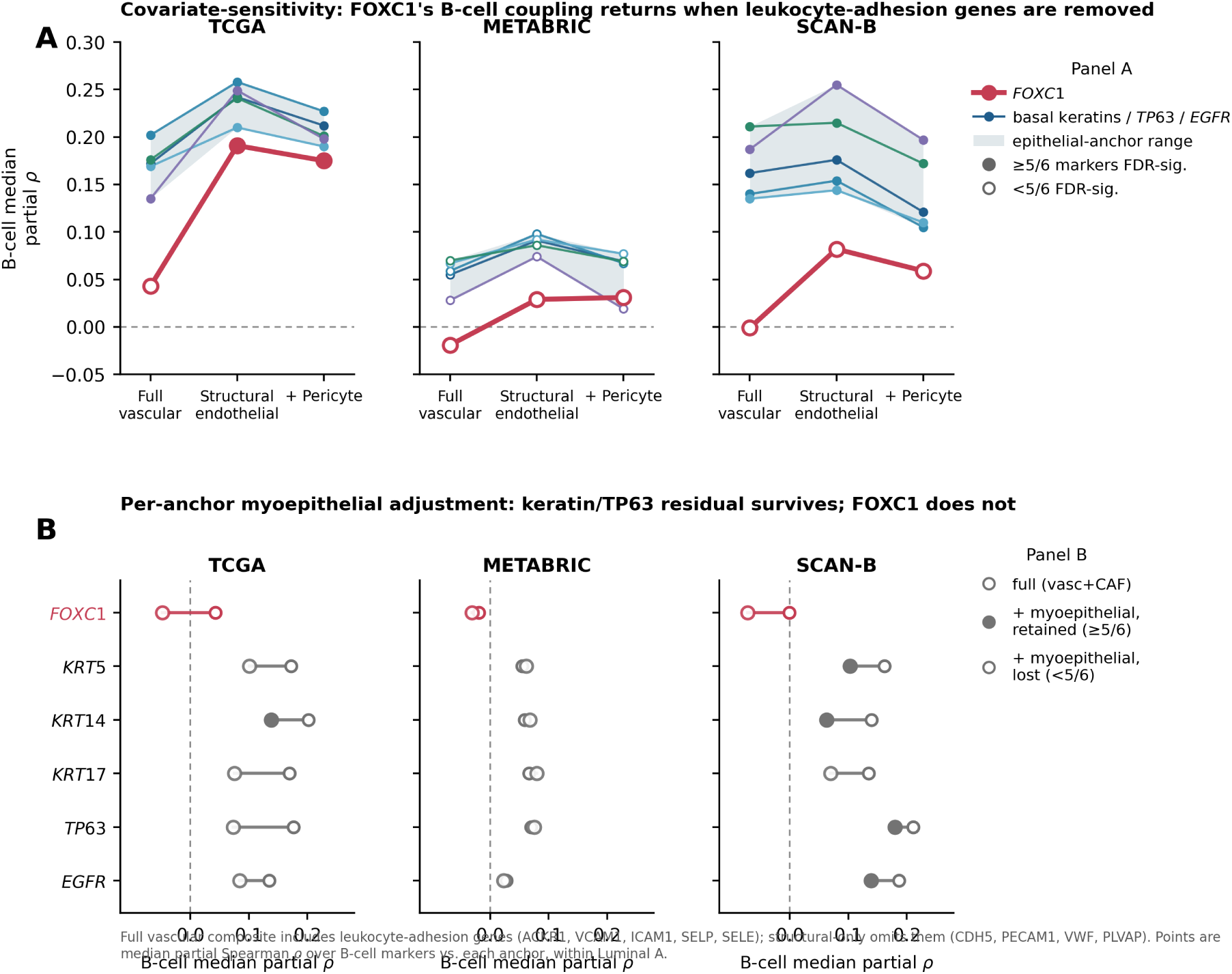
The *FOXC1* collapse is a compartment partition, not an over-adjustment artefact. (A) Covariate-sensitivity of *FOXC1*’s B-cell coupling across vascular-composite definitions in three cohorts: under the trafficking-inclusive vascular+CAF composite the coupling is near-zero, but under a structural-only endothelial composite (*CDH5*/*PECAM1*/*VWF*/*PLVAP*, dropping the leukocyte-adhesion genes *ACKR1*/*VCAM1*/*ICAM1*/*SELP*/*SELE*) it returns (TCGA 0.04 → 0.19; SCAN-B −0.00 → 0.08), localising the signal to adhesion-competent, immune-recruiting endothelium; adding pericyte markers changes little. (B) Per-anchor effect of additionally adjusting for a myoepithelial composite: *FOXC1*’s residual coupling moves negative, whereas keratin/*TP63* anchors retain B-cell coupling after the same adjustment (SCAN-B *KRT5*/*TP63*/*EGFR* remain FDR-significant). *FOXC1*’s signal thus partitions fully into non-malignant compartments (adhesion endothelium + myoepithelium); the epithelial anchors do not. Values in Supplementary Tables S9, S10.

### Benchmarking against alternative basal-lineage anchors FOXC1 collapses while basal cytokeratins and TP63 retain a compartment-independent B-cell and TLS coupling

To distinguish an anchor-specific effect from a framework-level negative result, we benchmarked five alternative basal-lineage anchor genes against *FOXC1* under identical vascular+CAF adjustment: *KRT5*, *KRT14*, *KRT17* (PAM50-defining basal cytokeratins), *TP63*, and *EGFR* (Fig. 5).

**Figure 5.**
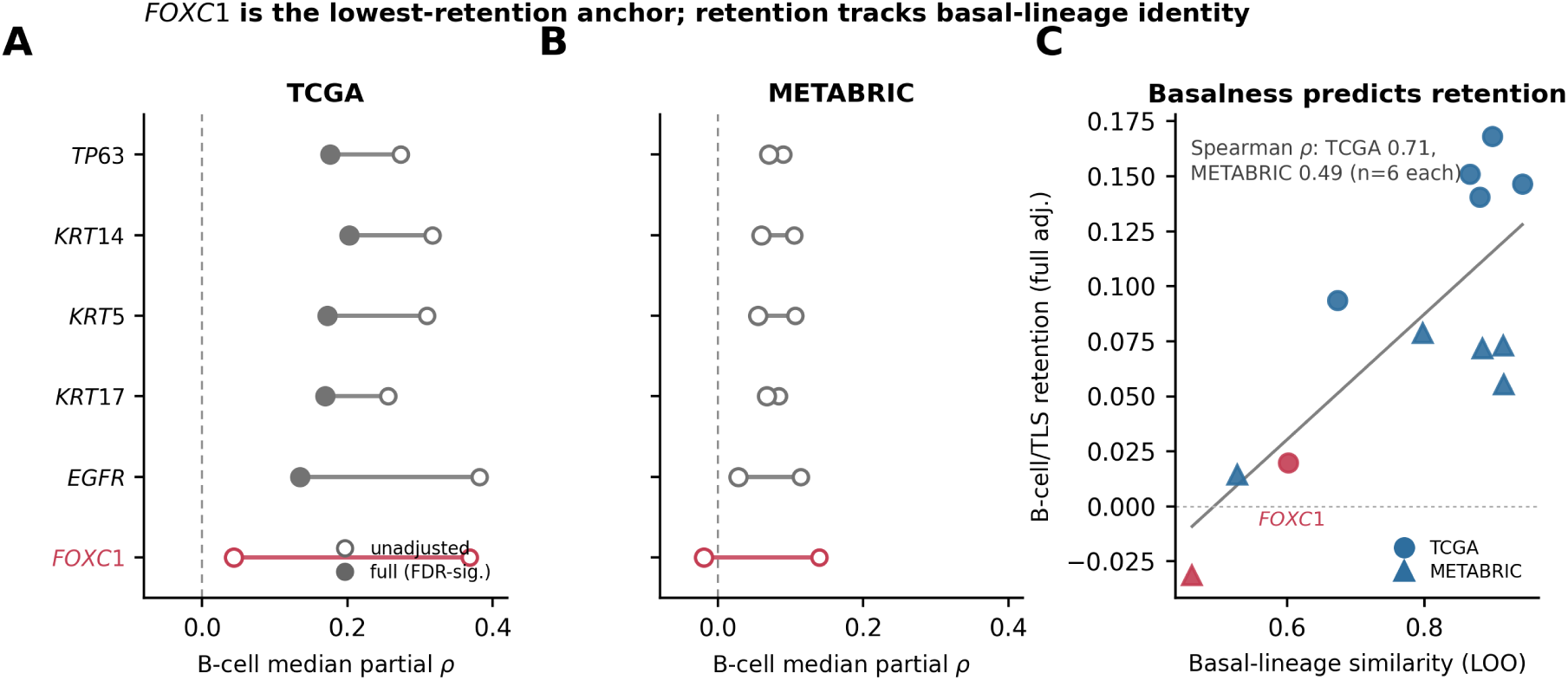
Which compartment carries the coupling distinguishes the anchors: *FOXC1*’s is vas-cular/myoepithelial, the keratins’ is epithelial-restricted. (A,B) B-cell median partial Spearman correlation, unadjusted → Full (Vascular+CAF), for six basal-lineage anchor genes in TCGA and METABRIC Luminal A. All six show comparable unadjusted couplings; under Full adjustment keratin/*TP63* anchors retain median *ρ* = 0.11–0.20 (6/6 B-cell markers FDR-significant in TCGA) while *FOXC1* collapses to *ρ* ≈ 0.04 (1/6). Open points, unadjusted; filled, Full (FDR-significant). (C) Retention under Full adjustment tracks basal-lineage identity: leave-one-out basal-lineage similarity versus B-cell/TLS retention across the six anchors (Spearman *ρ* = 0.71 TCGA, 0.49 METABRIC). *FOXC1*, the least basal and most vascular anchor, sits at the low-retention corner. Under a structural-only endothelial composite this distinction narrows (Fig. 4A), so the contrast is which compartment carries the coupling, not that *FOXC1* is uniquely fragile.

The pattern is anchor-specific and reproducible, though the retained residuals are small and, at the single-anchor level, not adjusted for myoepithelial content (Limitations). In TCGA, keratin/*TP63* anchors retain median partial Spearman *ρ* = 0.11–0.20 across B-cell and TLS-organising categories with 5/6 to 6/6 B-cell markers FDR-significant per anchor: B-cell median partial correlations were *ρ* = +0.20 (*KRT14*, 6/6 FDR-sig), +0.18 (*TP63*, 6/6), +0.17 (*KRT5*, 6/6), +0.17 (*KRT17*, 6/6), +0.14 (*EGFR*, 5/6), and +0.04 (*FOXC1*, 1/6); the same rank ordering held for TLS-organising chemokines. Averaged across B-cell and TLS, *FOXC1* sits last among the six with a mean residual ∼8-fold smaller than the top four keratin/*TP63* anchors. Both METABRIC and the SCAN-B RNA-seq cohort reproduce the same rank ordering: in SCAN-B all five keratin/*TP63* anchors retain the B-cell coupling (6/6 markers FDR-significant each), *TP63* again leads on B-cell+TLS retention (median partial *ρ* = +0.20) and *FOXC1* again ranks last of the six (*ρ* = −0.00, 0/6 B-cell markers FDR-significant), so *FOXC1* is the lowest-retention anchor in all three cohorts across two sequencing platforms (Supplementary Table S8). This ranking reflects the full vascular+CAF model; under structural-only endothelial adjustment *FOXC1*’s coupling returns to keratin-like magnitude (above). The anchors are therefore distinguished not by whether their coupling attenuates under a single adjustment but by which compartment carries it: *FOXC1*’s is endothelial and myoepithelial and is removed by adjusting for either, whereas the keratin/*TP63* residual is epithelial-restricted and survives vascular, CAF and myoepithelial adjustment (below).

Ranking each anchor by its correlation with a leave-one-out composite of the other basal-lineage genes – a measure of how faithfully each anchor tracks the tumour-cell basal programme in bulk expression – and against the vascular composite explains this mechanistically. In both cohorts *FOXC1* ranks last (6/6) by basal-lineage similarity (*ρ* = 0.60 in TCGA, 0.46 in METABRIC) and first (1/6) by vascular correlation (*ρ* = 0.67 in TCGA, 0.57 in METABRIC), while the four keratin/*TP63* anchors occupy the top four positions on both rankings in the opposite direction (basalness *ρ* = 0.87–0.94 in TCGA; vascularity *ρ* = 0.16–0.39) (Supplementary Table S5). Anchor basalness is positively associated with compartment-adjusted B-cell/TLS retention across the six anchors (Spearman *ρ* = 0.71 in TCGA, 0.49 in METABRIC; reported as descriptive given *n* = 6, with permutation and bootstrap uncertainty specified in Methods), and vascularity inversely so (*ρ* = −0.60 and −0.83). *FOXC1*’s compartment-adjusted collapse is therefore the direct quantitative consequence of *FOXC1* carrying the largest fraction of non-tumour-cell (vascular) signal in bulk Luminal A expression among the six candidate anchors tested; basal cytokeratins and *TP63*, being epithelial-restricted, carry a signal less confounded by vascular/CAF composition through the same adjustment; whether that retained signal is malignant-epithelial or reflects non-malignant myoepithelial content is examined directly below.

### A basal-epithelial lineage composite retains the residual signal, does not depend on PAM50-defining basal genes, and is robust to composite construction

We next asked whether a combined basal-lineage composite behaves like the individual keratin/*TP63* anchors. To operationalise the Centaur phenotype as a graded programme, we constructed a basal-epithelial lineage composite score from eleven tumour-cell-intrinsic basal markers (*KRT5*, *KRT14*, *KRT17*, *KRT6B*, *EGFR*, *TP63*, *CDH3*, *MIA*, *SFRP1*, *S100A2*, *FOXC1*) that deliberately excludes stromal, myoepithelial, and EMT genes. Under joint vascular+CAF adjustment, this composite retains a residual B-cell signature in TCGA (median partial *ρ* = +0.20; 6/6 B-cell markers FDR-significant), together with TLS-organising chemokines (*ρ* = +0.13; 3/5 genes), mature-DC markers (*ρ* = +0.20; 2/2 genes), and, more weakly, cytotoxic effectors (*ρ* = +0.06; 3/9 genes). METABRIC replicates the pattern with attenuated magnitudes (B-cell *ρ* = +0.05, 2/6 genes; TLS *ρ* = +0.08, 2/5 genes; mature-DC *ρ* = +0.12, 2/2 genes), while a third, independent RNA-seq cohort – SCAN-B (GSE96058, *n* = 1,540 LumA) – reproduces it at a magnitude closer to TCGA than to METABRIC (B-cell *ρ* = +0.17, 6/6 genes; TLS *ρ* = +0.09, 3/5 genes; mature-DC *ρ* = +0.13, 2/2 genes). The residual is positive and directionally consistent across all three cohorts – strong (6/6 B-cell markers FDR-significant) in the RNA-seq cohorts TCGA and SCAN-B, attenuated (2/6) in microarray METABRIC – consistent with a platform dynamic-range contribution, though cohort composition may also contribute (Fig. 2; Supplementary Tables S3, S8).

This residual composite–immune coupling is partially sensitive to myoepithelial content. Because basal cytokeratins and *TP63* are also canonical myoepithelial markers, we added a single-cell-derived myoepithelial-content score (Methods) to the vascular+CAF adjustment and recomputed the composite–immune partial correlations in the Luminal A strata. This attenuated the coupling in TCGA – median partial *ρ* across the B-cell, TLS and cytotoxic categories fell from 0.139 to 0.069, and FDR-significant immune genes from 12 to 8 – while leaving the weaker METABRIC association largely intact (0.056 to 0.048; six genes in both models). A residual signal therefore persists after myoepithelial adjustment, but a substantial fraction – roughly half in TCGA – of the bulk basal–immune coupling reflects variable myoepithelial content rather than malignant basal-programme activation (Supplementary Table S7). Because this signature rests on a single atlas, we treat the malignant-associated fraction as an upper bound and read the basal-lineage axis as a partly-myoepithelial, partly-malignant bulk signal rather than a purely tumour-cell one.

Applying the same myoepithelial adjustment to the single-gene anchors (Supplementary Table S10) attenuates but does not abolish the epithelial anchors’ residual: in SCAN-B, *KRT5*, *TP63* and *EGFR* remain 6/6 B-cell-FDR-significant after myoepithelial adjustment (median partial *ρ* 0.10–0.18) and *KRT14* 5/6, whereas *FOXC1* falls to a negative residual (*ρ* = −0.07). The keratin/*TP63* anchors thus retain a myoepithelial-adjusted, epithelial-restricted B-cell/TLS component that *FOXC1* entirely lacks – the operational distinction between the two at the single-anchor level.

Because four of the eleven genes in this composite (*KRT5*, *KRT14*, *KRT17*, *FOXC1*) are also PAM50-defining basal centroid markers, we tested whether the residual reflects a soft-gradient toward the PAM50 Basal centroid rather than a within-LumA lineage axis. An alternative composite constructed exclusively from non-PAM50 tumour-cell-intrinsic basal markers (*KRT6B*, *EGFR*, *TP63*, *CDH3*, *MIA*, *SFRP1*, *S100A2*) retained the residual B-cell coupling in TCGA at essentially unchanged magnitude and gene count (median partial *ρ* = +0.21 versus +0.20; 6/6 genes FDR-significant in both), together with equivalent TLS-organising and mature-DC signatures. The residual is therefore demonstrably not a re-encoding of PAM50 subtype-classifier gene overlap.

Leave-one-out sensitivity, rebuilding the composite eleven times with a different constituent excluded per iteration, confirmed robustness to single-gene removal: in both cohorts, no LOO iteration reversed the sign of the B-cell, TLS, or mature-DC residual (Supplementary Table S6). In TCGA, per-category median partial *ρ* varied by at most Δ*ρ* ≈ 0.03 for B cell (6/6 markers FDR-significant in every iteration), TLS, and mature-DC. The largest single-gene contribution came from *TP63*, consistent with its independent identification in the anchor benchmark as a top-performing basal anchor; *TP63* exclusion attenuated the B-cell, TLS, and mature-DC residuals by 5–12% but did not eliminate them. The cytotoxic residual, already the weakest of the four categories at *ρ* = 0.06 in TCGA, was substantially more *TP63*-dependent (attenuated by ∼80%); we do not carry the cytotoxic residual forward as a primary claim.

The signal also survives over-adjustment for FRC/TLS scaffolding covariates in TCGA: under vascular+CAF+FRC adjustment the B-cell partial correlation rises from *ρ* = +0.20 to *ρ* = +0.24 (6/6 FDR-significant), a statistical-suppression effect in which collinear FRC variance had partly masked the tumour-cell basal signal. In METABRIC, substituting an epithelial-minus-stromal purity proxy for the TCGA-only CPE covariate left the correlations essentially unchanged (B-cell *ρ* = 0.05 vs 0.04); because the proxy’s luminal-cytokeratin numerator (*KRT8/18/19*) sits opposite the composite’s basal cytokeratins, shared dependence would deflate rather than inflate the residual, so this stability rules out that circularity.

### The stromal-vascular remodelling co-occurs with a specific mature-vasculature signature

Beyond the compartment-adjustment characterisation above, the Centaur-associated vascular programme has a specific molecular signature. *FOXC1* correlated with a coordinated endothelial programme (*ACKR1*, *PECAM1*, *SELP*, *CDH5*) at effect sizes exceeding every immune-gene correlation, but *not* with *VEGFA* (mildly anti-correlated), indicating mature differentiated vasculature rather than the immature angiogenesis of a growing tumour (Supplementary Table S1); the covariate-sensitivity analysis (Results) locates *FOXC1*’s immune coupling specifically in the leukocyte-adhesion markers of this programme (*ACKR1*, *SELP*, *SELE*, *VCAM1*, *ICAM1*), identifying *FOXC1* as a marker of the immune-recruiting, adhesion-competent vessel wall; in parallel, fibroblastic reticular cell markers (*CXCL12*, *PDPN*, *IL7*) [27, 28] and cancer-associated fibroblast markers (*PDGFRA*, *ACTA2*, *PDGFRB*) were coordinately elevated, and xCell2 [29] deconvolution corroborated the gene-level pattern. Whether this vascular remodelling is driven by the Centaur tumour-cell state, is a substrate on which it develops, or shares an upstream cause with it, is not resolvable by bulk observational data (see Discussion).

### The basal-lineage axis is a tumour-population phenomenon, not within-cell co-expression

We next asked whether the basal-lineage axis reflects individual cancer cells co-expressing basal and luminal programmes, or graded activation across tumours resolved only in bulk. Two population-level observations establish the axis as a graded within-Luminal A phenomenon. First, dose-binning across the *FOXC1* range (Supplementary Fig. S1) showed the basal-lineage composite gained +1.23 SD in TCGA and +1.02 SD in METABRIC from Q1 to Q4, while the luminal-identity composite (*ESR1*, *FOXA1*, *GATA3*) declined by only −0.49 SD and −0.32 SD; basal gain exceeded luminal attenuation ∼2.5–3-fold, without accompanying proliferative gain (Supplementary Fig. S3), so basal-lineage identity is acquired across the axis without basal-like proliferative kinetics. Second, PAM50 subtype-call stability under Gaussian-noise perturbation (200 iterations per cohort) [24] confirmed the axis is a within-Luminal A phenomenon and not a boundary-classification artefact (median LumA classification 1.00 in both Centaur-high and Centaur-low subgroups; mean 0.995 vs 0.970 in METABRIC, 0.999 vs 0.993 in TCGA; Supplementary Fig. S4).

At single-cell resolution, however, the axis does not resolve into individual dual-lineage cells. In ER^+^ cancer cells (*n* = 11,878; GSE176078 [34]) the basal-lineage anchors were expressed in only a small minority (*KRT5* 0.4%, *KRT17* 1.8%, *TP63* 0.1%), and their co-detection with *ESR1* did not survive adjustment for sequencing depth (depth-adjusted odds ratios *KRT5* 1.24, *p* = 0.55; *KRT14* 1.01, *p* = 0.95; *KRT17* 0.97, *p* = 0.87; *TP63* 7.73, *p* = 0.054 on 12 cells), and anchor^+^ cells showed no elevation of luminal-identity score over anchor^−^ cells (Wilcoxon *p >* 0.2 for all anchors). *FOXC1* gave the highest raw co-detection with *ESR1* (depth-adjusted OR 2.20, *p* = 0.027), and – unlike the keratin/*TP63* anchors – this OR persists rather than collapses under DecontX ambient-RNA correction (decontaminated OR 2.36, *p* = 0.021; Methods), so it is not attributable to endothelial ambient spillover. It rests, however, on the ∼0.5% of ER^+^ malignant cells in which *FOXC1* is detected at all, is therefore underpowered, and concerns *FOXC1* – the gene here reclassified as vascular – rather than a basal-lineage anchor, so it does not bear on the within-cell status of the basal-lineage axis (Fig. 3C). We therefore interpret the basal-lineage axis as a tumour-population property rather than a within-cell lineage-hybrid state; the present single-cell data, limited by the low per-cell capture of these markers, neither support within-cell hybridity nor exclude a rare hybrid subpopulation, and we do not claim one.

### The Centaur axis is orthogonal to clinical genomic tools and distinct from established immune signatures

The molecular phenotype replicated in METABRIC: nineteen of thirty-seven immune genes reached FDR *<* 0.05, led by the same TLS-organising chemokines, B-cell markers, and cytotoxic effectors; the stromal programme replicated 22/25 genes; and proliferation-independence held (Supplementary Table S1).

Against proliferation-based clinical tools, the axis is orthogonal: *FOXC1* shows near-zero correlation with Oncotype DX and MammaPrint proxies and with the proliferation gene *AURKA*, while correlating with *ESR1* and the PAM50 subtype-of-recurrence component (Table 3; Fig. 6A). Two Luminal A tumours with identical Oncotype DX or MammaPrint proxy scores may differ markedly in Centaur status. Against established immune signatures, *FOXC1* itself correlated only weakly with the Tumor Inflammation, Cabrita TLS, and Helmink / Petitprez B-cell signatures (*ρ* ≤ 0.29); a secondary composite built from the *FOXC1*-coordinated immune/stromal genes overlapped these signatures more strongly (*ρ* = 0.69–0.78) but retained an association with survival after signature adjustment (Table 2), indicating that the phenotype’s coupling to lineage hybridity is not a restatement of any published immune signature.

**Figure 6.**
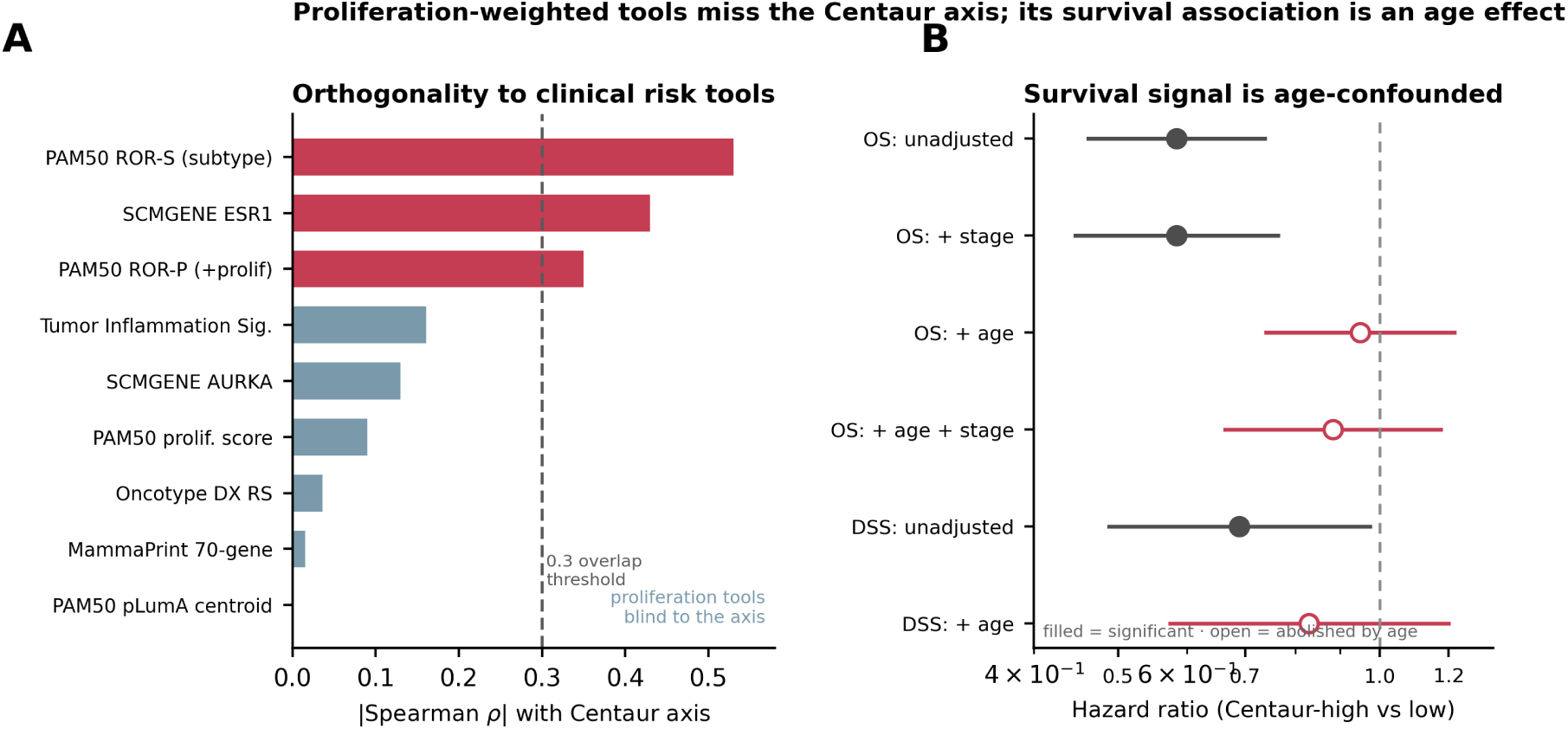
The Centaur axis is orthogonal to proliferation-based clinical tools, and its survival association is an age effect. (A) Absolute Spearman correlation of the Centaur axis with established genomic risk scores in Luminal A; proliferation-weighted tools (Oncotype DX, MammaPrint, PAM50 proliferation) fall well below the 0.3 substantial-overlap threshold (dashed line) and are blind to the axis, whereas lineage-based scores (PAM50 ROR-S, SCMGENE *ESR1*) exceed it as expected for a basal–luminal axis (full values in Table 3). (B) METABRIC overall- and disease-specific-survival hazard ratios for Centaur-high versus -low across adjustment models: the unadjusted association (OS HR = 0.58 (95% CI 0.46–0.74)) survives stage adjustment but is abolished by age adjustment (OS age-adjusted HR = 0.95 (95% CI 0.74–1.22), *p* = 0.68; DSS age-adjusted HR = 0.83 (95% CI 0.57–1.20), *p* = 0.32), identifying the survival signal as largely an age effect. Filled points, significant; open, abolished by age.

**Table 2:** Cox proportional-hazards regression summary: age-confounding of the Centaur survival association. All models used two-sided tests. HR *<* 1 indicates better survival in the Centaur-high group. The primary Centaur definition is the basal-epithelial lineage axis (Methods). The critical comparison is between the stage-adjusted rows and the age-adjusted rows: METABRIC OS association is retained under stage adjustment but abolished under age adjustment, and the pattern replicates on disease-specific survival. Sequential mediation, single-gene *FOXC1* replication, and signature-adjusted models in Supplementary Table S2. TCGA discovery rows rest on 21 events, are hypothesis-generating, and do not survive joint age+stage adjustment.

| Cohort | Endpoint | Predictor / Model | *n* | Events | HR | 95% CI | *p* |
| --- | --- | --- | --- | --- | --- | --- | --- |
| *TCGA-BRCA: discovery (hypothesis-generating, 21 events)* |  |  |  |  |  |  |  |
| TCGA | OS | *FOXC1*-high vs low (no radiation) | 218 | 21 | 0.28 | 0.09–0.84 | 0.016 |
| TCGA | OS | *FOXC1* + age + stage (no radiation) | 218 | 21 | 0.42 | — | 0.145 |
| *METABRIC: Centaur phenotype (basal-epithelial lineage axis); primary definition* |  |  |  |  |  |  |  |
| METABRIC | OS | Centaur-high vs low, unadjusted | 700 | 294 | 0.584 | 0.462–0.737 | $6.2 \times 10^{-6}$ |
| METABRIC | OS | + stage | 700 | 294 | 0.584 | 0.446–0.764 | $9.0 \times 10^{-5}$ |
| METABRIC | OS | + age | 700 | 294 | **0.949** | **0.740–1.218** | **0.68** |
| METABRIC | OS | + age + stage | 700 | 294 | 0.883 | 0.664–1.175 | 0.39 |
| *METABRIC: disease-specific survival* |  |  |  |  |  |  |  |
| METABRIC | DSS | Centaur-high vs low, unadjusted | 699 | 131 | 0.689 | 0.488–0.974 | 0.035 |
| METABRIC | DSS | + age | 699 | 131 | **0.829** | **0.574–1.199** | **0.32** |

**Table 3:** Orthogonality of the Centaur axis to established clinical genomic risk tools (TCGA LumA, *n* = 571). Spearman *ρ* between *FOXC1* expression and each clinical tool score or component. The 0.3 threshold (dashed line in Fig. 6A) marks substantial overlap; all values fall below this threshold, indicating that proliferation-weighted tools do not capture the Centaur axis. *ESR1* and ROR-S show moderate negative correlations, consistent with the Centaur axis being a lineage-axis (luminal–basal) rather than proliferation-axis phenomenon.

| Tool / Score | Type | Spearman $\rho$ | Interpretation |
| --- | --- | --- | --- |
| Oncotype DX recurrence score | Proliferation-weighted | +0.036 | Near-null; blind to Centaur axis |
| MammaPrint (70-gene) | Proliferation-weighted | −0.015 | Near-null; blind to Centaur axis |
| SCMGENE *AURKA* component | Proliferation | −0.130 | Near-null; proliferation axis orthogonal |
| SCMGENE *ESR1* component | Luminal identity | −0.430 | Moderate–strong; Centaur varies on lineage axis |
| PAM50 ROR-S (subtype only) | Subtype/lineage | −0.530 | Moderate–strong; captures basal–luminal component |
| PAM50 ROR-P (+ prolifer.) | Subtype + proliferation | −0.350 | Attenuated by proliferation weighting |
| PAM50 proliferation score | Proliferation | −0.090 | Near-null; confirms proliferation independence |
| PAM50 pLumA centroid | Subtype centroid | $\approx 0.000$ | Null; restricted range within LumA |
| *Immune and TLS signature comparators* |  |  |  |
| Tumor Inflammation Sig. (TIS) | Immune/pan-T | 0.161 | *FOXC1* weakly correlated (composite $\rho = 0.69$); composite retains OS after TIS adjustment (HR = 0.71, $p < 0.001$) |
| Cabrita TLS (12-gene) | TLS/B-cell | 0.260 | *FOXC1* modestly correlated (composite $\rho = 0.73$); composite retains OS after Cabrita adjustment (HR = 0.81, $p = 0.011$) |
| Helmink B-cell (9-gene) | B-cell/TLS | 0.264 | *FOXC1* modestly correlated (composite $\rho = 0.76$); composite retains OS after Helmink adjustment (HR = 0.80, $p = 0.008$) |
| Petitprez B-cell (10-gene) | B-cell/TLS | 0.291 | Highest composite overlap ($\rho = 0.78$); borderline after adjustment (HR = 0.85, $p = 0.065$) |
| Poudel stem-like subtype | Heterocellular | $\approx 0.200$ | Weak; only 47% of Centaur is stem-like |
| Poudel transit-amplifying | Heterocellular (proliferative) | $\approx -0.300$ | Centaur depleted in TA subtype |

*Note.* Oncotype DX and MammaPrint scores were not directly measured in TCGA; values shown are based on published correlation analyses with TCGA RNA-seq data using the validated computational implementations of each signature [38, 39]. ROR scores were computed using the PAM50 centroid-based method [1]. Poudel subtype assignments were obtained from the published per-sample classifications for the TCGA cohort [37].

### The apparent survival advantage is largely explained by patient age

Finally, we examined whether the Centaur phenotype carries independent prognostic weight, and here the evidence is clear but cautionary. Defining the Centaur phenotype by the basal-epithelial lineage axis, Centaur-high Luminal A tumours in METABRIC showed a strong unadjusted association with improved overall survival (HR = 0.58 (95% CI 0.46–0.74); *p <* 10^−5^; Fig. 6B; Table 2), present in both radiation-treated and radiation-näıve strata, with no radiation interaction (Wald *p* = 0.82). However, this association was almost entirely explained by patient age. Adjusting for tumour stage left the effect intact (HR = 0.58), but adjusting for age abolished it (HR = 0.95, *p* = 0.68); the asymmetry is the signature of age confounding rather than an independent tumour-biological effect. The same pattern held on disease-specific survival, the endpoint least susceptible to competing-cause mortality: the unadjusted association (HR = 0.69, *p* = 0.035) did not survive age adjustment (HR = 0.83, *p* = 0.32), though we note this endpoint is underpowered at 131 events and the age-adjusted point estimate remains below unity. Consistent with this, the basal-lineage axis was enriched in younger patients, and age was a dominant predictor in every adjusted model.

The discovery-cohort TCGA data showed a directionally similar but even more fragile picture: in the radiation-näıve subgroup, high *FOXC1* was associated with improved overall survival (Table 2), but this subgroup split is non-randomised – radiation status reflects treatment received rather than random assignment, so it is subject to confounding by indication – and rested on only 21 deaths, so the estimate is correspondingly imprecise and likely inflated, and it did not survive joint adjustment for age and stage; we therefore present it as supplementary and confirmatory only, and base the survival conclusion on the better-powered METABRIC analysis above. Across both cohorts, therefore, the Centaur phenotype does not provide survival information beyond standard clinical variables, and its apparent prognostic signal is largely an age effect. We emphasise that this does not diminish the molecular phenotype – which is robust and reproducible – but it does mean the Centaur axis should be understood as a biological rather than a prognostic entity on present evidence, with any clinical prognostic role requiring prospective, adequately powered, covariate-controlled evaluation.

## Discussion

This study makes two positive contributions to the interpretation of Luminal A biology. First, single-cell and compartment-resolved analysis localises bulk *FOXC1* to the tumour vasculature and reassigns its apparent adaptive-immune coupling to adhesion-competent, immune-recruiting endothelium rather than to the malignant compartment. Second, compartment-adjusted basal cytokeratins and *TP63* recover a genuine, proliferation-independent basal-lineage axis, the Centaur axis, within Luminal A tumour cells. More broadly, the analysis shows that a bulk transcriptomic biomarker can be dominated by a minority stromal compartment, so compartment-aware adjustment is needed before a bulk correlation is read as tumour-cell biology. We discuss each below, together with the interpretive limits of a compartment-level reanalysis.

### The Centaur phenotype and its interpretive framing

The Centaur phenotype is a continuous lineage-hybrid axis within PAM50 Luminal A that is proliferation-independent, orthogonal to proliferation-based clinical tools, and cuts across categorical heterocellular subtype boundaries (only 47% of Centaur tumours were classified as stem-like by Poudel et al. [37]; the association with heterocellular classification was driven by depletion in the transit-amplifying subtype, consistent with the Centaur axis marking a less proliferative, more differentiated state than the transit-amplifying compartment).

Beyond the compartment-adjustment, single-cell-attribution and anchor-benchmark evidence that constitutes this paper’s findings, the coordinated programme (Supplementary Fig. S5) admits two speculative readings, offered as an interpretive frame rather than a result. A cancer-cell-autonomous reading has basal-lineage-anchored tumour cells elevating myoepithelial paracrine factors (*FGF2*, *NRG1*, *HGF*) that seed FRC-like CAF activation and TLS scaffolding (*CCL19*/*CCL21*) supporting lymphocyte recruitment [30, 31]; a vascular/stromal-first reading has a coordinated vascular/stromal expansion producing both the bulk signal and the coupled recruitment. For *FOXC1* specifically, single-cell attribution settles this in favour of the vascular-first reading (90% vessel-wall, 1.8% malignant); what remains observational is the direction of any coupling between the tumour-cell basal-lineage axis and the immune programme, which requires spatial and functional work.

### FOXC1 in bulk LumA is principally a stromal-vascular marker; the compartment-adjusted framework recovers the residual tumour-cell signal

Stated plainly, the central finding is not that *FOXC1* is a poor marker but that bulk *FOXC1* in Luminal A primarily reflects stromal and vascular biology, while compartment-adjusted basal cytokeratins and *TP63* more faithfully track the residual tumour-linked basal-lineage programme – itself only partly malignant (above). The principal analytical contribution is a purity- and stroma-adjusted reanalysis of the *FOXC1*-associated basal-lineage programme in Luminal A bulk data, now corroborated at single-cell resolution (above). Three points warrant emphasis in the Discussion beyond their statement in Results. First, the anchor benchmark demonstrates that the compartment-adjusted collapse of *FOXC1* is not a framework failure indicting the basal-lineage programme itself; alternative basal anchors (*KRT5*, *KRT14*, *KRT17*, *TP63*) retain the tumour-cell coupling to B-cell and TLS-organising signatures under identical adjustment, and *FOXC1*’s collapse follows quantitatively from its being the least basal-like of the six candidate anchors tested. Second, the framework has a candidate diagnostic property, suggestive across the six anchors tested rather than formally powered: anchor basalness (leave-one-out similarity to the other candidates) is positively associated with compartment-adjusted retention (Spearman *ρ* = 0.71 in TCGA; descriptive, *n* = 6), and the *FOXC1*-specific result is one consequence of that broader relationship rather than a special property of *FOXC1*. Within the anchor benchmark, *FOXC1* (90% vessel-wall attribution) functions as an internal negative control for the tumour-cell channel and the basal cytokeratins (0.4–1.8% malignant single-cell detection but epithelial-restricted bulk expression) as internal positive controls; the framework’s operating characteristics on genes with ground-truth compartment attribution outside the basal-lineage panel remain to be established. Third, the framework is a candidate template for other bulk expression settings in which a tumour-cell driver has documented stromal or vascular activity, rather than a benchmarked general-purpose method; calibration against positive- and negative-control genes and against existing deconvolution-adjustment approaches would be a natural next step.

On transparency: this is a computational reanalysis, and neither the *FOXC1*-centred nor the basal-keratin-anchored framing was pre-registered; what was specified in advance was the compartment-adjustment framework, whose diagnostic property (basalness predicts retention) holds independently of which anchor is the a priori focus.

### Biological implications and clinical potential

Whether the compositional distinction between Centaur and non-Centaur Luminal A has clinical conse-quences – for endocrine-therapy response, recurrence pattern, or long-term outcome – cannot be established from observational bulk data and we do not claim it here. Two Luminal A tumours with identical Oncotype DX or MammaPrint proxy scores may nonetheless differ substantially in Centaur status; whether this lineage-classifier-blind axis carries clinical utility would require prospective, treatment-annotated evaluation. We note that several features of the phenotype – proliferation independence, enrichment in younger patients, and the absence of an independent adverse prognostic signal – are collectively consistent with, though they do not establish, a relatively indolent biological character; this would be one natural hypothesis for prospective testing. Compartment-resolving spatial methods – when they include *FOXC1* (current commercial breast panels either lack *FOXC1* or offer whole-transcriptome resolution only at multi-cell resolution) – would allow the compartment attribution above to be tested directly.

### Limitations

This study is observational; all associations are correlational. Two scope limits bear on the central claim. First, the single-cell attribution establishes the cellular *source* of *FOXC1* transcripts, not the *function* of the residual malignant pool: reclassifying bulk *FOXC1* as principally stromal-vascular concerns the origin of the bulk signal, not the significance of low-abundance malignant *FOXC1*, which transcription factors can retain at modest expression and which prior perturbation studies [17, 15] probed mechanistically; resolving it requires perturbational data. Second, the vascular composite includes adhesion/trafficking genes (*ACKR1*, *VCAM1*, *ICAM1*, *SELP*, *SELE*) that mark immune-recruiting endothelium rather than pure confounders; the structural-only endothelial, pericyte, and per-anchor myoepithelial specifications (Results; Supplementary Tables S9, S10) localise *FOXC1*’s coupling to the adhesion-competent vasculature and confirm the keratin/*TP63* residual is epithelial-restricted. Deconvolution-based specifications (e.g. CIBERSORTx, EPIC) were not tested, and the myoepithelial signature rests on a single atlas (GSE176078).

The compartment-adjustment and anchor-benchmark results hold in three cohorts across two sequencing platforms, but the survival, dose-binning, PAM50-stability, and leave-one-out analyses were run only in TCGA and METABRIC (SCAN-B’s deposition lacks purity, stage, and outcome fields), and TCGA radiation status reflects treatment received rather than random assignment. At single-cell resolution the basal-lineage axis is a tumour-population rather than a within-cell property, and the data do not resolve whether the coupled immune programme is downstream of tumour-cell basal activation or of coordinated stromal/vascular expansion. One specific alternative is that basal cytokeratins and *TP63*, canonical myoepithelial markers detected in only 0.1–1.8% of ER^+^ malignant cells (Fig. 3C), partly reflect variable retained myoepithelial content; adjusting for a single-cell-derived myoepithelial score removes roughly half of the bulk basal–immune coupling in TCGA (Results), leaving a residual whose definitive attribution would require multiplexed in-situ staining. The basalness–retention relationship is descriptive (*n* = 6). The Centaur composite was derived from TCGA-internal correlations and evaluated in METABRIC without formal cross-validation, mitigated by its reproduction in the independent SCAN-B cohort and by a PAM50-independent composite recovering it at essentially unchanged magnitude (Results). Oncotype DX/MammaPrint orthogonality used RNA-seq-derived proxies rather than clinically reported assay scores, and single-cell anchor co-detection is limited by low per-cell capture. Establishing *FOXC1*’s in-situ vascular localisation, the direction of the basal–immune coupling, and any clinical utility requires spatial (with *FOXC1* coverage), functional, and prospective validation.

## Materials and methods

### Cohorts

We analysed three Luminal A (LumA) cohorts on two platforms: TCGA-BRCA RNA-seq (DESeq2-normalised; *n* = 571 Luminal A; GDC), METABRIC microarray (*n* = 700 Luminal A; cBioPortal `brca metabric`), and SCAN-B RNA-seq (log_2_-FPKM; *n* = 1,540 Luminal A; GSE96058 [35], an independent sequencing pipeline). PAM50 calls and clinical data were taken from each source’s published annotation; analyses were restricted to PAM50-assigned Luminal A samples. *ACKR1* was absent from the SCAN-B deposition (its vascular composite used the remaining eight genes), which lacks purity/stage/DSS/treatment fields and so was used for correlation and anchor analyses only. Full cohort processing is in Supplementary Methods.

### Centaur (basal-lineage) composite

The Centaur phenotype was defined by a basal-epithelial lineage score: the mean of per-gene column-wise z-scored log_2_ expression across eleven tumour-cell-intrinsic basal markers (*KRT5*, *KRT14*, *KRT17*, *KRT6B*, *EGFR*, *TP63*, *CDH3*, *MIA*, *SFRP1*, *S100A2*, *FOXC1*), with equal (not data-driven) weighting; per-gene z-scoring precedes averaging, and myoepithelial/EMT/stromal markers were deliberately excluded so the score reflects tumour-cell basal activation rather than stromal content. Centaur-high/low groups were a within-Luminal A median split. Leave-one-out sensitivity (Supplementary Table S6; Supplementary Methods) confirms no single constituent drives the residual.

### Compartment-adjusted partial correlations

To separate tumour-cell-intrinsic effects from stromal-compositional co-variation, we computed partial Spearman correlations (`ppcor::pcor.test`) between the axis of interest (single-gene *FOXC1* or the basal composite) and each immune gene, adjusting for compartment covariates: a vascular composite (*CDH5*, *PECAM1*, *VWF*, *ACKR1*, *SELP*, *SELE*, *PLVAP*, *VCAM1*, *ICAM1*), a CAF composite (*PDGFRA*,

*PDGFRB*, *ACTA2*), and (in TCGA only) CPE tumour purity [36]. Five nested models were fit per gene per cohort (unadjusted; purity; vascular; CAF; full vascular+CAF), with two-sided *p* and Benjamini–Hochberg FDR within cohort × axis × model, and attenuation quantified as median *ρ*_full_*/ρ*_unadjusted_ per immune category. Because malignant cells contribute only 1.8% of *FOXC1* transcripts (Fig. 3), the vascular composite cannot mediate a tumour-cell *FOXC1* effect, so the confounder assumption is independently validated for *FOXC1*; for the keratin/*TP63* anchors it is not, and partial correlations alone cannot exclude mediation there. Two sensitivity analyses localised the *FOXC1* signal (Supplementary Methods): a *structural-only* endothelial composite (*CDH5*/*PECAM1*/*VWF*/*PLVAP*) omitting leukocyte-adhesion genes (*ACKR1*/*VCAM1*/*ICAM1*/*SELP*/*SELE*), ± a pericyte composite (Supplementary Table S9); and a single-cell-derived 40-gene *myoepithelial* composite added to the adjustment set (Supplementary Tables S7, S10).

### Anchor benchmarking

The partial-correlation analysis was repeated with each of five alternative single-gene basal anchors (*KRT5*, *KRT14*, *KRT17*, *TP63*, *EGFR*) replacing *FOXC1* under identical adjustment; anchors were scored by per-category median partial *ρ* and FDR-significant gene count under full adjustment. A PAM50-independent composite and additional covariate checks (FRC/TLS, purity-proxy, compartment-adjusted proliferation) are in Supplementary Methods.

### Single-cell analysis

GSE176078 [34] annotations were mapped to six compartments. Per gene we computed compartment transcript-share, per-compartment detection rate, and per-patient malignant:endothelial ratios (≥20 cells/compartment). Co-detection of each anchor with *ESR1* in ER^+^ malignant cells was tested by Fisher exact and depth-adjusted logistic regression, and ambient-RNA sensitivity assessed with DecontX (celda) on raw counts. Luminal-identity and proliferation definitions and full procedures are in Supplementary Methods.

### Survival and statistics

OS and DSS were evaluated by Kaplan–Meier and Cox regression in METABRIC (censored 180 months) and OS in TCGA (censored 4,000 days, radiation-stratified), comparing unadjusted with age- and stage-adjusted models. Published immune/TLS signatures [41, 40, 32, 33] and xCell2 deconvolution [29] are described in Supplementary Methods. All *p*-values are two-sided (Benjamini–Hochberg FDR within panel); analyses used R 4.3.0. Code: https://github.com/danielyehoshua123/Centaur-phenotype-analysis.

## Acknowledgements

We thank the developers of xCell2 for making their tool publicly available, and the TCGA, METABRIC, and GSE176078 data-generating consortia for making their datasets publicly available.

## Competing interests

The authors declare that no competing interests exist.

## Funding

This work was supported by the Israel Science Foundation and the Technion – Israel Institute of Technology. The funders had no role in study design, data collection and interpretation, or the decision to submit the work for publication.

## Author contributions

D.E. Yehoshua: Conceptualization; Methodology (initial); Investigation (initial); Formal analysis (initial); Writing – original draft (initial). J.C. Bingham: Conceptualization; Methodology; Software; Investigation; Formal analysis; Validation; Visualization; Data curation; Writing – original draft; Writing – review & editing; Project administration.

## Data availability

All datasets analysed in this study are publicly available. TCGA-BRCA data are available through the GDC portal (https://portal.gdc.cancer.gov); METABRIC data through cBioPortal (https://www.cbioportal.org/study/summary?id=brca_metabric); and the GSE176078 single-cell RNA-seq dataset through GEO (https://www.ncbi.nlm.nih.gov/geo/query/acc.cgi?acc=GSE176078). Analysis code is available at https://github.com/danielyehoshua123/Centaur-phenotype-analysis.

*Supplementary Table S1: cross-platform replication of FOXC1 immune and stromal correlations in TCGA and METABRIC Luminal A (per-gene Spearman ρ, FDR, and replication disposition). Provided as supplementary CSV*.

*Supplementary Table S2: extended Cox proportional-hazards results. Sequential mediation rows (TCGA), single-gene FOXC1 non-replication (METABRIC), and immune/stromal composite adjusted for published signatures (TIS, Cabrita TLS, Helmink B-cell, Petitprez). Extends Table 2 in the main text*.

*Supplementary Table S3: per-gene partial Spearman correlations of FOXC1 and the basal-epithelial lineage composite with immune genes under five nested compartment-adjustment models in TCGA and METABRIC. Provided as supplementary CSV (`results/01 partial correlations.csv`)*.

*Supplementary Table S4: per-anchor, per-immune-category median partial Spearman correlations for the six basal-lineage anchor genes benchmarked in Fig. 5, under unadjusted and joint vascular+CAF-adjusted models in TCGA and METABRIC. Provided as supplementary CSV (`results/07B alternative anchors.csv`)*.

*Supplementary Table S5: per-anchor Spearman correlation with a leave-one-out composite of the other basal-lineage genes (’basalness’), and with the vascular composite (’vascularity’), in TCGA and METABRIC Luminal A. FOXC1 ranks lowest in basalness and highest in vascularity of the six anchors tested, matching its position last in the compartment-adjusted retention ranking of Fig. 5. Provided as supplementary CSV (`results/09 anchor ranking.csv`)*.

*Supplementary Table S6: leave-one-out sensitivity of the basal-epithelial lineage composite. For each of the eleven constituent genes, the composite was rebuilt excluding that gene and vascular+CAF-adjusted partial Spearman correlations with immune genes recomputed. Table reports, per cohort per immune category, the reference (all-eleven-gene) median partial ρ and FDR-significant gene count, and the range across the eleven leave-one-out iterations. Provided as supplementary CSV (`results/10 composite LOO summary` with per-iteration per-gene detail in `results/10 composite LOO sensitivity.csv`)*.

*Supplementary Table S7: myoepithelial-content sensitivity of the basal-epithelial lineage composite. A 40-gene myoepithelial signature (top panel: genes derived from GSE176078 myoepithelial cells, excluding basal anchors, pan-epithelial, vascular, CAF, smooth-muscle/contractile and technical modules) was scored per tumour and added to the vascular*+*CAF(+purity) adjustment. The bottom panel reports, per cohort and per immune category, the median partial Spearman ρ and FDR-significant gene count under the vascular*+*CAF model versus the vascular*+*CAF*+*myoepithelial model in Luminal A TCGA and METABRIC. The basal–immune coupling attenuates by roughly half in TCGA (median ρ* 0.139 → 0.069; 12 → 8 *genes) and minimally in METABRIC (*0.056 → 0.048*; six genes), indicating that a substantial part of the bulk basal signal reflects myoepithelial content while a residual persists. Provided as supplementary CSVs (`results/myoepithelial signature.txt`, `results/myoepithelial adjustment.csv`)*.

**Supplementary Table S8.** SCAN-B (GSE96058) third-cohort validation, LumA *n* = 1,540. Median partial Spearman correlations within the SCAN-B Luminal A stratum under joint vascular+CAF adjustment, computed identically to the TCGA and METABRIC analyses.

| Axis | Category | Median $\rho$ | FDR-sig | Genes |
| --- | --- | --- | --- | --- |
| *FOXC1* (unadj) | B cell | +0.287 | 6 | 6 |
| *FOXC1* (unadj) | TLS | +0.280 | 5 | 5 |
| *FOXC1* (unadj) | Mature DC | +0.253 | 2 | 2 |
| *FOXC1* (unadj) | Cytotoxic | +0.280 | 9 | 9 |
| *FOXC1* (full adj) | B cell | -0.001 | 0 | 6 |
| *FOXC1* (full adj) | TLS | -0.093 | 1 | 5 |
| *FOXC1* (full adj) | Mature DC | -0.061 | 0 | 2 |
| *FOXC1* (full adj) | Cytotoxic | -0.085 | 0 | 9 |
| Composite (full adj) | B cell | +0.165 | 6 | 6 |
| Composite (full adj) | TLS | +0.086 | 3 | 5 |
| Composite (full adj) | Mature DC | +0.131 | 2 | 2 |
| Composite (full adj) | Cytotoxic | +0.037 | 2 | 9 |

| Anchor | B-cell median $\rho$ | FDR-sig | B-cell+TLS median $\rho$ |
| --- | --- | --- | --- |
| *TP63* | +0.211 | 6 | +0.202 |
| *EGFR* | +0.187 | 6 | +0.172 |
| *KRT5* | +0.162 | 6 | +0.148 |
| *KRT14* | +0.140 | 6 | +0.128 |
| *KRT17* | +0.135 | 6 | +0.123 |
| *FOXC1* | −0.001 | 0 | −0.007 |

**Supplementary Table S9.** Covariate-specification sensitivity of the *FOXC1* collapse (B-cell category, full-adjusted). Median partial Spearman *ρ* (FDR-significant/6 markers) under three vascular specifications: M0, the primary vascular+CAF composite; M1, structural-only endothelial (*CDH5*/*PECAM1*/*VWF*/*PLVAP*)+CAF, dropping the leukocyte-adhesion genes; M2, M1 + pericyte composite. *FOXC1*’s coupling returns under M1/M2 while the keratin/*TP63* range is stable, showing the collapse is carried by the adhesion markers.

| Cohort | Model | *FOXC1* $\rho$ (FDR/6) | keratin/*TP63/EGFR* $\rho$ range |
| --- | --- | --- | --- |
| TCGA | M0 vasc+CAF | +0.04 (1/6) | +0.14 to +0.20 (5–6/6) |
| TCGA | M1 struct-endo+CAF | +0.19 (6/6) | +0.21 to +0.26 (6/6) |
| TCGA | M2 struct+peri+CAF | +0.18 (6/6) | +0.19 to +0.23 (6/6) |
| SCAN-B | M0 vasc+CAF | −0.00 (0/6) | +0.14 to +0.21 (6/6) |
| SCAN-B | M1 struct-endo+CAF | +0.08 (4/6) | +0.14 to +0.26 (6/6) |
| SCAN-B | M2 struct+peri+CAF | +0.06 (3/6) | +0.11 to +0.20 (6/6) |
| METABRIC | M0 vasc+CAF | −0.02 (0/6) | +0.03 to +0.07 (1–3/6) |
| METABRIC | M1 struct-endo+CAF | +0.03 (1/6) | +0.07 to +0.10 (3/6) |
| METABRIC | M2 struct+peri+CAF | +0.03 (1/6) | +0.02 to +0.08 (1–2/6) |

**Supplementary Table S10.** Per-anchor myoepithelial adjustment (B-cell category). Median partial Spearman *ρ* (FDR-significant/6) for each anchor under the vascular+CAF (“full”) model versus that model plus the 40-gene myoepithelial-content covariate (“+myo”), in TCGA and SCAN-B. Keratin/*TP63* anchors retain FDR-significant B-cell coupling after myoepithelial adjustment (6/6 for *KRT5*, *TP63*, *EGFR* in SCAN-B), whereas *FOXC1* falls to a negative residual.

| Anchor | TCGA |  |  | SCAN-B |  |  |
| --- | --- | --- | --- | --- | --- | --- |
| | full | +myo | $\Delta\rho$ | full | +myo | $\Delta\rho$ |
| *FOXC1* | +0.04 (1/6) | −0.05 (0/6) | −0.09 | −0.00 (0/6) | −0.07 (0/6) | −0.07 |
| *KRT5* | +0.17 (6/6) | +0.10 (3/6) | −0.07 | +0.16 (6/6) | +0.10 (6/6) | −0.06 |
| *KRT14* | +0.20 (6/6) | +0.14 (5/6) | −0.06 | +0.14 (6/6) | +0.06 (5/6) | −0.08 |
| *KRT17* | +0.17 (6/6) | +0.08 (2/6) | −0.09 | +0.14 (6/6) | +0.07 (3/6) | −0.07 |
| *TP63* | +0.18 (6/6) | +0.07 (3/6) | −0.10 | +0.21 (6/6) | +0.18 (6/6) | −0.03 |
| *EGFR* | +0.14 (5/6) | +0.08 (3/6) | −0.05 | +0.19 (6/6) | +0.14 (6/6) | −0.05 |

## Declaration of generative AI and AI-assisted technologies in the writing process

During the preparation of this work the authors used large-language-model AI assistants (Claude, Anthropic) to support code drafting, iterative manuscript editing, and analytical discussion. After using these tools the authors reviewed and edited the content as needed and take full responsibility for the content of the publication. All analyses, statistical results, interpretations, and scientific conclusions are the authors’ own and were independently verified against source data and against the code provided in the Code availability statement.

## Supplementary Methods

### TCGA-BRCA data

RNA-seq data (DESeq2-normalised counts) from TCGA-BRCA (*n* = 1, 099 tumours; *n* = 571 Luminal A) were obtained from the GDC portal. PAM50 subtypes were taken from published assignments. All analyses were restricted to PAM50-assigned samples with matched clinical data.

### METABRIC validation data

METABRIC expression data (Illumina microarray log_2_ intensities; *n* = 1, 980 total; *n* = 700 Luminal A) and clinical data were downloaded from cBioPortal (`brca metabric`). PAM50 subtypes were taken from the `CLAUDIN SUBTYPE` column. Survival analysis used overall survival (OS) and disease-specific survival (DSS; “Died of Disease” as the event), censored at 180 months.

### SCAN-B validation data

SCAN-B RNA-seq expression data (log_2_-FPKM; *n* = 1,540 Luminal A within the series) and sample annotation were obtained from Gene Expression Omnibus series GSE96058 [35], an independent, population-based RNA-seq cohort profiled on a sequencing pipeline distinct from TCGA. PAM50 subtype calls and overall survival were taken from the series sample metadata. Compartment-adjustment and anchor-benchmark analyses were run identically to the discovery and first-validation cohorts, using the same gene sets, composite construction, and vascular+CAF adjustment (the vascular marker *ACKR1* was absent from the SCAN-B deposition, so its vascular composite used the remaining eight genes); because the GEO deposition does not include tumour purity, stage, DSS, or treatment fields, SCAN-B was used for the correlation and anchor analyses and not for survival modelling.

### Expression analyses

*FOXCUT*–*FOXC1* correlations across subtypes were computed by Spearman *ρ*. Gene-level immune, stromal, and paracrine panel correlations were computed within the Luminal A stratum by Spearman *ρ* with two-sided *p*-values and Benjamini–Hochberg FDR correction applied to each panel separately.

### Centaur phenotype definition: basal-epithelial lineage axis

The Centaur phenotype was defined by a basal-epithelial lineage score: the mean of column-wise per-gene z-scored log_2_ expression values across eleven tumour-cell-intrinsic basal markers (*KRT5*, *KRT14*, *KRT17*, *KRT6B*, *EGFR*, *TP63*, *CDH3*, *MIA*, *SFRP1*, *S100A2*, *FOXC1*). We chose equal weighting across genes rather than data-driven weights (e.g. from PCA or regression against an outcome) so that the composite reflects a biologically-motivated basal-lineage definition rather than a cohort-specific fit; leave-one-out sensitivity (Results, Supplementary Table S6) confirms the residual signal is not driven by any single constituent, and the anchor benchmark (Fig. 5) confirms that the compartment-adjusted result is recoverable through any of the individual keratin/*TP63* anchors. Per-gene z-scoring before averaging prevents any single high-variance keratin from dominating the score. Myoepithelial, EMT, and stromal markers (*ACTA2*, *MYH11*, *CALD1*, *CNN1*, *VIM*, *NGFR*) were deliberately excluded so that the score reflects tumour-cell basal-lineage activation rather than stromal content, avoiding mechanical inflation of the downstream stromal and immune associations. Centaur-high and Centaur-low groups were defined by a median split of this score within the Luminal A stratum. The basal-lineage axis correlated with *FOXC1* alone (*ρ* = 0.75 pan-cohort) but was not identical to it; within Luminal A, *FOXC1* correlated with the basal programme with *FOXC1* itself excluded (*ρ* = 0.46).

### Secondary immune/stromal composite (signature benchmarking)

For the immune-signature benchmarking analysis only, a separate immune/stromal composite score was computed as the mean of column-wise z-scored expression across nine genes showing the strongest *FOXC1* correlations among the curated immune/stromal panels (*CCL21*, *CCL19*, *CD79B*, *ACKR1*, *PECAM1*, *PRF1*, *CD8A*, *PDGFRA*, *CXCL12*). This composite is distinct from the basal-lineage axis used to define the Centaur phenotype and was used to characterise the relationship between the *FOXC1*-coordinated immune programme and established immune/TLS signatures. No survival data were used in gene selection for either score.

### Compartment-adjusted partial correlations

To distinguish tumour-cell-intrinsic effects from stromal-compositional co-variation in bulk expression data, we computed partial Spearman correlations between the axis of interest (single-gene *FOXC1* or the basal-epithelial lineage composite) and each immune gene, adjusted for compartment covariates. A vascular composite was defined as the mean of column-wise z-scored expression across *CDH5*, *PECAM1*, *VWF*, *ACKR1*, *SELP*, *SELE*, *PLVAP*, *VCAM1*, and *ICAM1*. A CAF composite was defined analogously across *PDGFRA*, *PDGFRB*, and *ACTA2*. For TCGA, tumour purity was estimated using the Aran et al. consensus purity estimate [36] (CPE); as CPE is not available for METABRIC, purity-adjusted models were computed in TCGA only. Five nested models were computed per gene per cohort: unadjusted, purity-adjusted (TCGA only), vascular-adjusted, CAF-adjusted, and jointly adjusted for vascular and CAF composites (“full” model). Partial Spearman correlations were computed using `ppcor::pcor.test` with Spearman method; complete cases only; two-sided *p*-values with Benjamini–Hochberg FDR correction applied within cohort, axis, and model strata. Attenuation was quantified as median *ρ*_full_*/ρ*_unadjusted_ per immune category within each cohort. This framework treats the vascular and CAF composites as confounders of the axis–immune association. For *FOXC1* this assumption is independently validated rather than merely assumed: because malignant cells contribute only 1.8% of *FOXC1* transcripts in ER^+^ tumours (Fig. 3A–C), the vascular composite cannot plausibly act as a downstream mediator of a tumour-cell *FOXC1* effect. For the basal cytokeratin and *TP63* anchors the confounder assumption is not independently established by single-cell data; partial correlations alone cannot distinguish confounding from mediation should tumour-cell basal activation causally elicit the vascular/CAF programmes, and we treat those anchor results accordingly.

#### Myoepithelial-content sensitivity

Because basal cytokeratins and *TP63* are canonical markers of normal myoepithelial cells, we tested whether the basal composite’s residual immune coupling could reflect variable retained myoepithelial content rather than malignant basal activation. A myoepithelial signature was derived from the GSE176078 atlas as the genes most specific to annotated myoepithelial cells, ranked by the difference between their mean expression in myoepithelial cells and their highest mean across the other compartments (malignant-epithelial, endothelial, perivascular and CAF), then excluding the basal anchors, pan-epithelial markers, the vascular and CAF composite genes, and smooth-muscle/contractile and technical (metallothionein, ribosomal, heat-shock, immediate-early, acute-phase) modules so that the score is independent of the vascular and CAF covariates already in the model and of ambient signal (40 genes; Supplementary Table S7). The signature was scored per tumour as a z-scored composite (as for the vascular and CAF composites) and added to the vascular+CAF (and, in TCGA, purity) adjustment set; the basal-composite–immune partial correlations were then recomputed under this expanded model in the Luminal A strata of TCGA and METABRIC using `ppcor::pcor.test`. The same myoepithelial covariate was added to each single-gene anchor model across all three cohorts to quantify the myoepithelial-attributable fraction of each anchor’s residual (Supplementary Table S10).

#### Covariate-specification sensitivity

To distinguish confounding from over-adjustment for endothelial mediators of immune recruitment, the *FOXC1* and anchor partial correlations were refit in TCGA, METABRIC and SCAN-B under (i) a structural-only endothelial composite (*CDH5*, *PECAM1*, *VWF*, *PLVAP*) omitting the leukocyte-adhesion/trafficking genes (*ACKR1*, *VCAM1*, *ICAM1*, *SELP*, *SELE*) present in the primary vascular composite, and (ii) that composite plus a pericyte composite (*RGS5*, *MCAM*, *NOTCH3*, *PDGFRB*, *KCNJ8*, *CSPG4*), retaining the CAF composite and (in TCGA) purity throughout (Supplementary Table S9).

### Anchor benchmarking and robustness analyses

To distinguish anchor-specific effects from framework-level results, the partial-correlation analysis was repeated with each of five alternative single-gene basal-lineage anchors – *KRT5*, *KRT14*, *KRT17*, *TP63*, and *EGFR* – in place of *FOXC1* as the axis of interest, in both cohorts under identical vascular+CAF adjustment. Anchor performance was assessed by per-category median partial Spearman correlation and count of FDR-significant genes under the full-adjustment model.

To assess robustness of the basal-epithelial lineage composite to inclusion of PAM50-defining basal genes, an alternative composite was constructed from tumour-cell-intrinsic basal markers excluded from the PAM50 basal centroid list (*KRT6B*, *EGFR*, *TP63*, *CDH3*, *MIA*, *SFRP1*, *S100A2*) and the partial-correlation analysis was rerun with this alternative composite as the axis of interest. As additional robustness checks: the adjustment model was extended in TCGA to include a fibroblastic reticular cell / TLS-scaffolding composite (*CXCL12*, *PDPN*, *IL7*, *LTB*, *LTA*, *TNFSF13B*) in addition to vascular+CAF; in METABRIC, an epithelial-minus-stromal purity proxy was computed as the mean-z-score difference between epithelial markers (*KRT8*, *KRT18*, *KRT19*, *EPCAM*, *CDH1*) and stromal markers (*COL1A1*, *COL1A2*, *COL3A1*, *VIM*) and used as an additional covariate in place of the CPE score available only in TCGA; and the proliferation analysis of Supplementary Fig. S3 was repeated under the same vascular+CAF adjustment as the immune analysis to assess whether proliferation-independence holds after compartment control.

### Composite construction sensitivity

To assess whether the compartment-adjusted immune-correlation results depend on any single constituent of the basal-epithelial lineage composite, we rebuilt the composite excluding each of its eleven constituent genes in turn (*KRT5*, *KRT14*, *KRT17*, *KRT6B*, *EGFR*, *TP63*, *CDH3*, *MIA*, *SFRP1*, *S100A2*, *FOXC1*) and repeated the vascular+CAF-adjusted partial-correlation analysis with each reduced composite as the axis of interest. Per-cohort per-category median partial Spearman correlations were compared across the eleven leave-one-out iterations against the reference (all-eleven-gene) composite (Supplementary Table S6); the residual is reported as robust to single-gene removal if no leave-one-out iteration reverses the sign of the reference residual in either cohort and no single-gene exclusion accounts for more than a small fraction of the reference median partial *ρ*.

### Immune/TLS signature scoring

Published immune signatures were scored as the mean of per-gene z-scored expression within the Luminal A stratum: the 18-gene Tumor Inflammation Signature [41], the Cabrita 12-gene TLS signature [40], the Helmink 9-gene B-cell signature [32], and the Petitprez B-cell proxy [33]. Correlations with *FOXC1* and the immune/stromal composite were computed by Spearman *ρ*; independent prognostic value was assessed by Cox models including the composite and each signature jointly.

### Survival analysis

Overall survival (OS) and disease-specific survival (DSS) were evaluated with Kaplan–Meier curves and Cox proportional-hazards regression in METABRIC (censored at 180 months) and OS in TCGA (censored at 4,000 days, stratified by radiation status). For each analysis, unadjusted models were compared with models adjusted for patient age and tumour stage, entered as z-scored and numeric covariates respectively; the divergence between stage-adjusted and age-adjusted estimates was used to assess confounding. The Centaur × radiation interaction was assessed by Wald test. Continuous-axis models were used alongside median-split models as more stable estimators at low event counts.

### Deconvolution

Cell-type proportion estimation used xCell2 [29] with the pan-cancer reference on TPM-normalised expression data. Spearman correlations between xCell2-derived cell-type scores and *FOXC1* were computed across LumA tumours.

### Lineage identity characterisation

Basal- and luminal-identity programmes were characterised using curated gene sets from established PAM50 identity markers, visualised as the basal–luminal composite plane (Fig. 1B) and quantified by *FOXC1*-quartile dose-binning (Supplementary Fig. S1). Per-marker differential expression between Centaur and non-Centaur Luminal A was tested by Wilcoxon rank-sum with Benjamini–Hochberg correction within each identity panel.

### Single-cell analysis

The GSE176078 scRNA-seq dataset [34] was used for single-cell analysis; cell-type annotations were taken directly from the Wu et al. metadata and mapped to six compartments (malignant epithelial, endothelial, perivascular, cancer-associated fibroblast, lymphoid, myeloid), with normal-epithelial cells excluded. For compartment attribution we computed, per gene, its total-contribution share – the fraction of all UMIs for that gene contributed by each compartment – together with per-patient malignant/endothelial expression ratios (patients with ≥20 cells in both compartments) and per-compartment detection rates. Profile concordance was the Pearson correlation between a gene’s six-compartment share vector and those of the vascular (*PECAM1*/*CDH5*/*VWF*) and epithelial (*EPCAM*) reference markers. Luminal A-like tumours were defined as the lowest-proliferation half of ER^+^ tumours, ranked by a per-tumour pseudobulk proliferation signature. For within-cell hybridity, co-detection of each basal anchor (*FOXC1*, *KRT5*, *KRT14*, *KRT17*, *TP63*) with *ESR1* among ER^+^ malignant cells was tested by Fisher exact test and by logistic regression controlling for log sequencing depth; luminal identity (*ESR1*/*FOXA1*/*GATA3* composite) of anchor^+^ versus anchor^−^ cells was compared by Wilcoxon test. Ambient-RNA sensitivity of the compartment attribution and of the *FOXC1*–*ESR1* co-detection was assessed with DecontX (celda) run on the raw GSE176078 counts using the provided cell-type labels as clusters; compartment shares, detection rates, and the depth-adjusted co-detection odds ratio were recomputed on the decontaminated counts and compared with the raw values.

### Tumour purity

Tumour purity was estimated using the Aran et al. consensus purity estimate [36] (CPE); 564 of 571 LumA tumours were matched. The radiation-näıve survival association was retained after purity adjustment (HR: 0.30 → 0.31).

### Statistical testing

All *p*-values are two-sided. Multiple testing correction used Benjamini–Hochberg FDR within each gene panel. All analyses were performed in R 4.3.0. Code is available at https://github.com/danielyehoshua123/Centaur-phenotype-analysis.

## Supplementary Figure legends

**Supplementary Figure S1.**
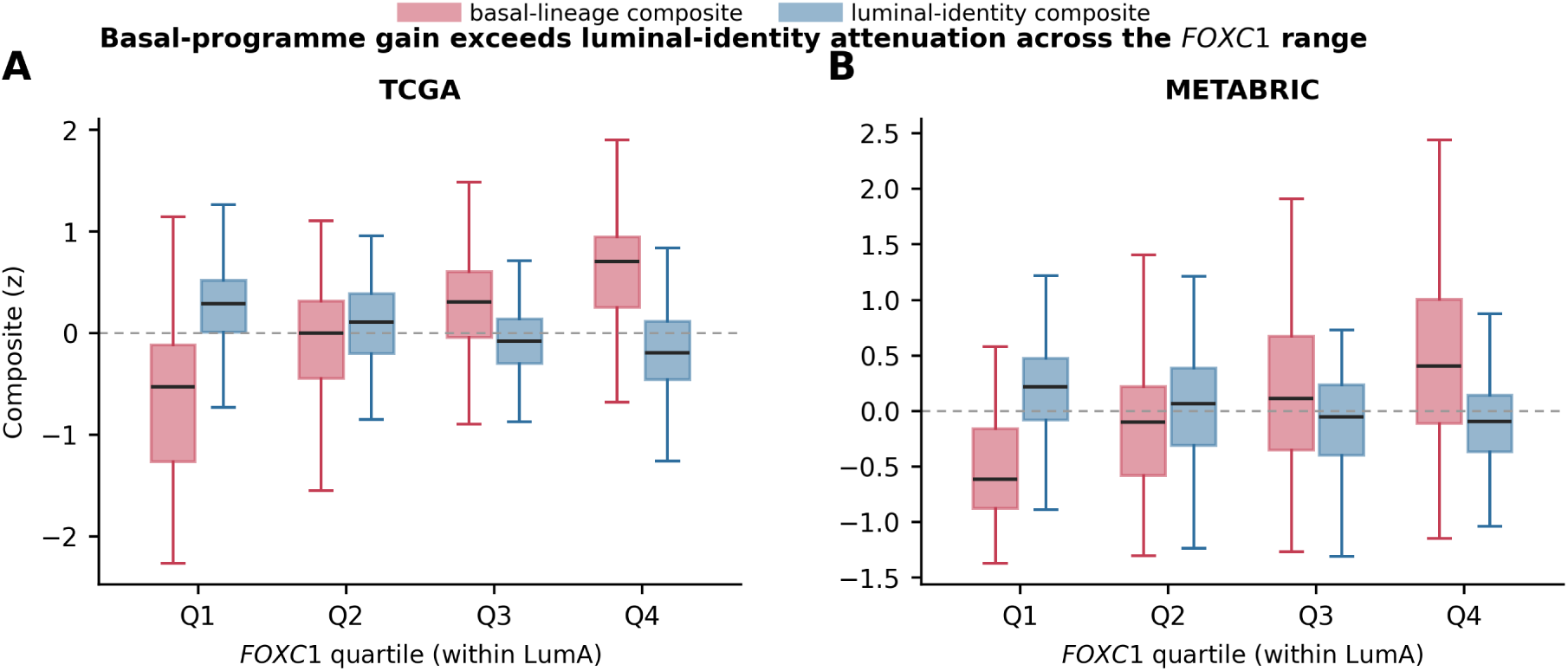
Basal-programme gain exceeds luminal-identity attenuation across the *FOXC1* range. Basal-lineage composite (excluding *FOXC1*) and luminal-identity composite for each Luminal A tumour, grouped by *FOXC1* quartile within the LumA stratum in TCGA (*n* = 571) and METABRIC (*n* = 700); boxes show median and IQR. The basal composite rises monotonically Q1→Q4 in both cohorts (adjacent-quartile Wilcoxon *p <* 10^−3^), while the luminal composite declines at roughly one-third the magnitude.

**Supplementary Figure S2.**
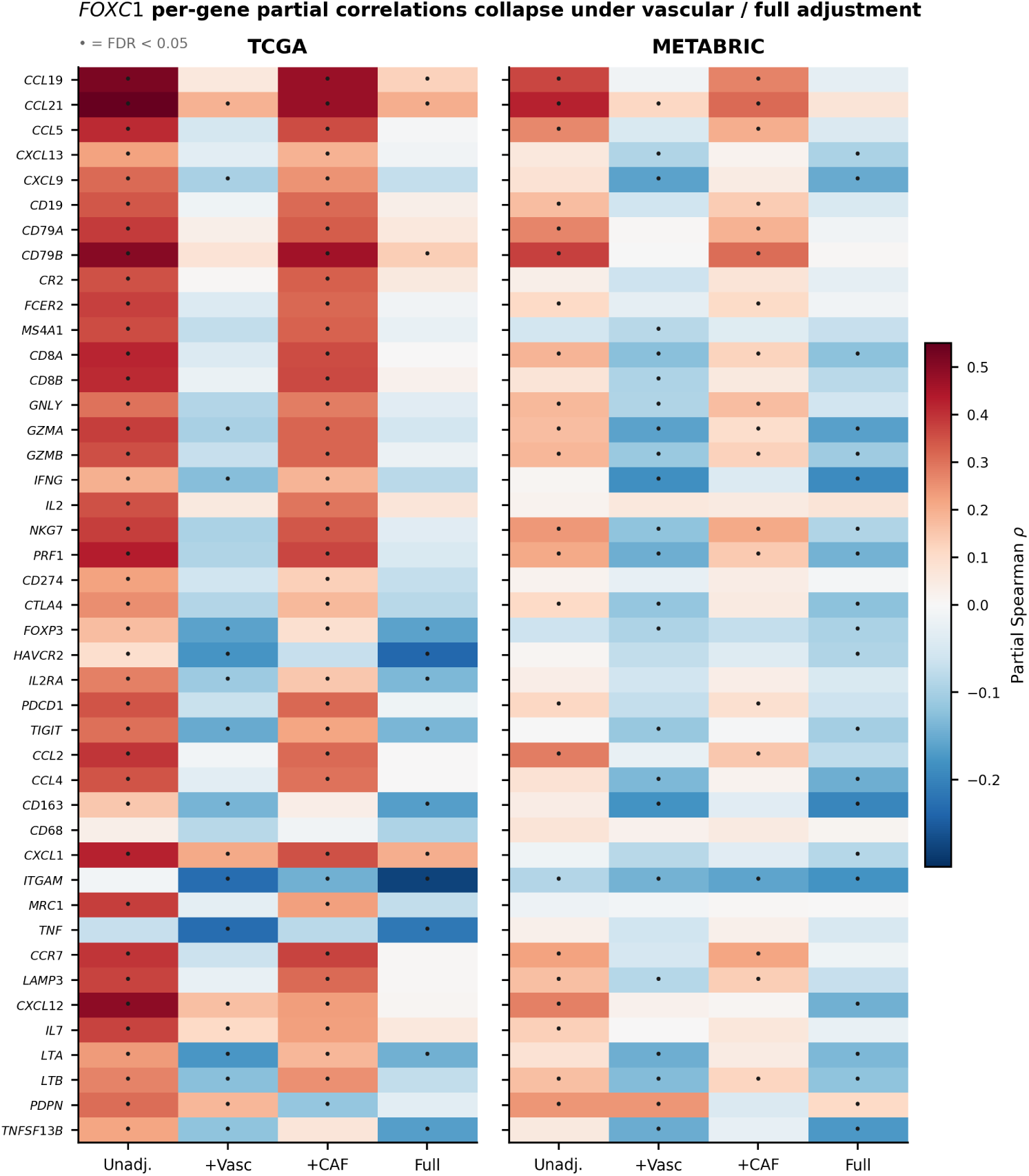
Per-gene *FOXC1* partial correlations across compartment-adjustment models. Partial Spearman correlations of *FOXC1* with each immune/stromal gene (rows, grouped by functional category) under the four nested models (Unadjusted, +Vascular, +CAF, Full) in TCGA and METABRIC Luminal A. Dots mark FDR*<* 0.05. The near-uniform collapse under vascular and Full adjustment underlies the category-level trajectories in Fig. 2.

**Supplementary Figure S3.**
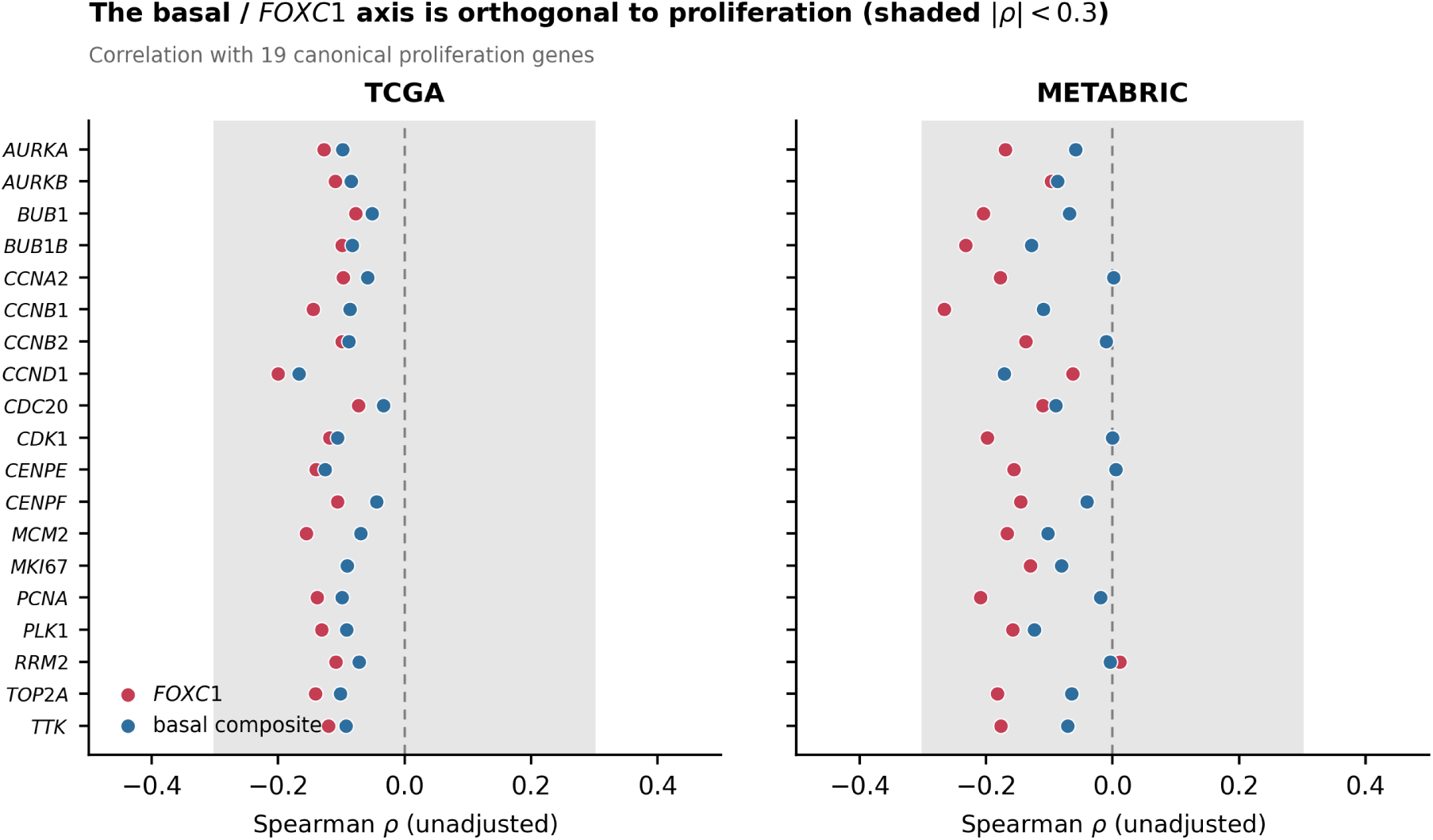
The basal-lineage axis is orthogonal to proliferation. Spearman correlations of *FOXC1* and the basal-lineage composite with 19 canonical proliferation genes (unadjusted) in TCGA and METABRIC Luminal A; nearly all fall within |*ρ*| *<* 0.3 (shaded), confirming the Centaur axis is not a proliferation signal.

**Supplementary Figure S4.**
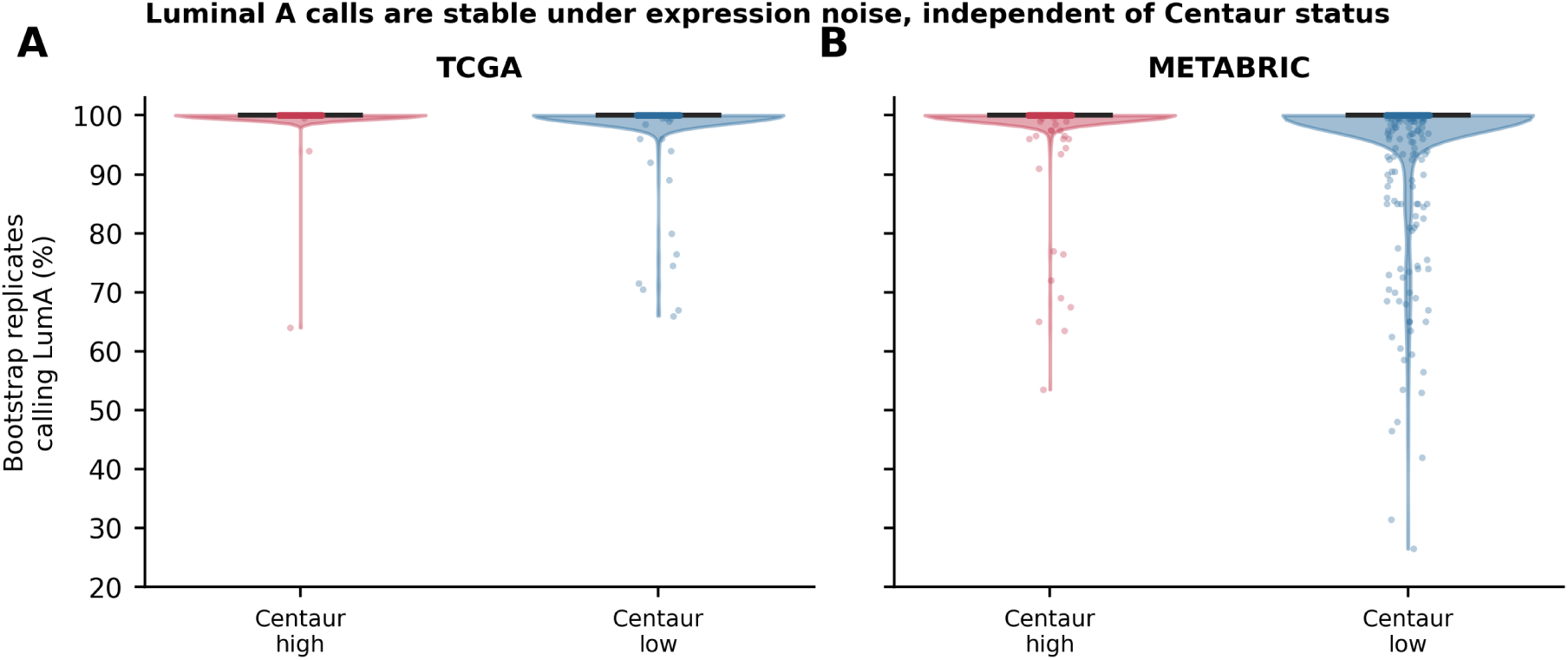
PAM50 subtype-call stability under expression-noise perturbation. For each baseline-Luminal A tumour, PAM50 subtype calls were recomputed across bootstrap iterations of Gaussian expression-noise perturbation. LumA retention is near-complete in every group (median 1.00; means 0.995 Centaur-high / 0.970 Centaur-low in METABRIC, 0.999 / 0.993 in TCGA), independent of Centaur status; Centaur-LumA tumours are not boundary-classified LumA cases.

**Supplementary Figure S5.**
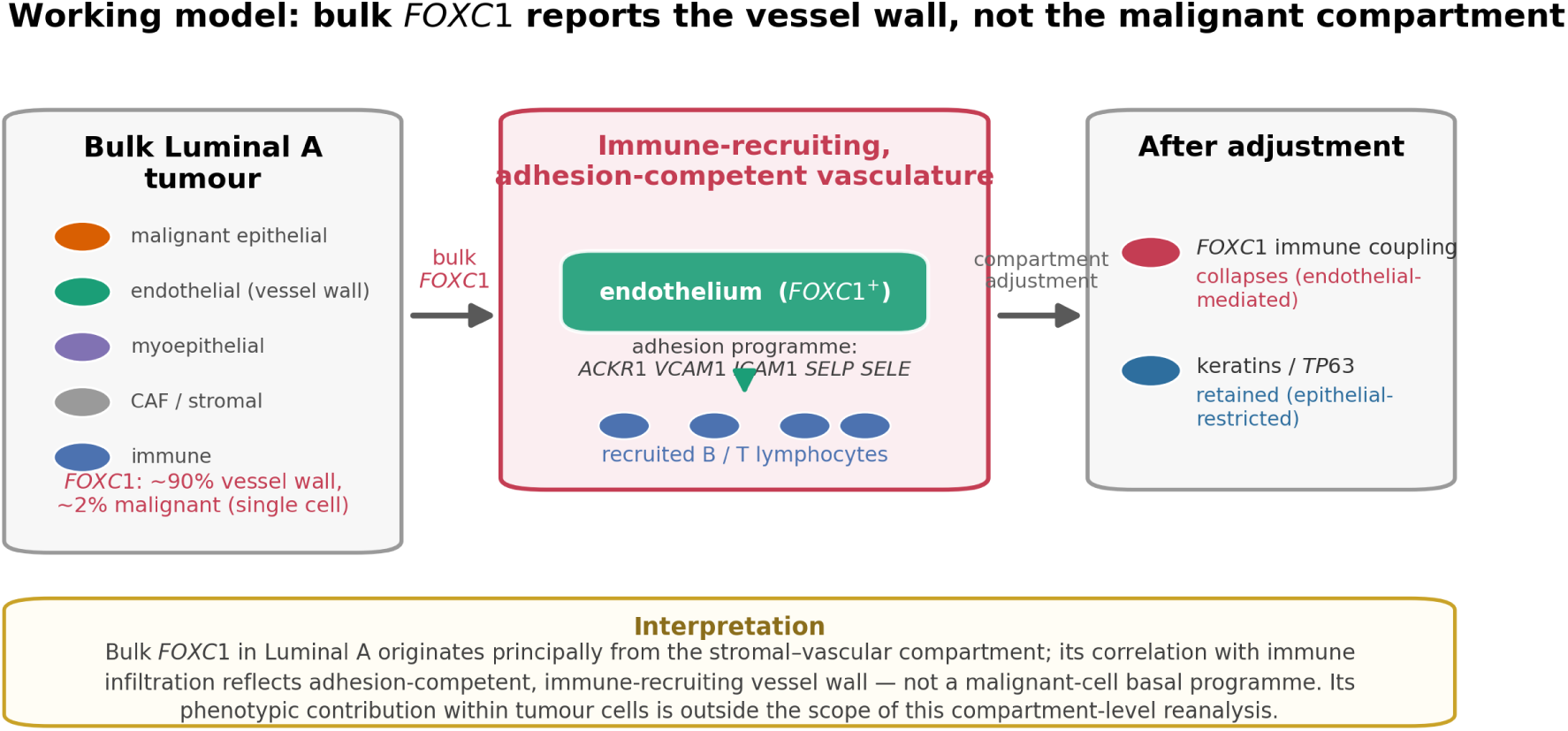
Working model: bulk *FOXC1* reports the vessel wall, not the malignant compartment. In Luminal A, ∼90% of bulk *FOXC1* derives from the vessel wall and ∼2% from malignant cells (single cell). The *FOXC1*-expressing endothelium is adhesion-competent and immune-recruiting (*ACKR1*/*VCAM1*/*ICAM1*/*SELP*/*SELE*), so *FOXC1*’s bulk correlation with lymphocyte infiltration is endothelial-mediated and is removed by compartment adjustment, whereas epithelial-restricted keratins/*TP63* are retained. Bulk *FOXC1* in Luminal A therefore originates principally from the stromal–vascular compartment; its phenotypic contribution within tumour cells is outside the scope of this compartment-level reanalysis. The directionality of any coupling between the epithelial basal-lineage axis and the coordinated immune programme is not resolved by observational bulk data (Discussion).

## Notes

https://portal.gdc.cancer.gov

https://www.cbioportal.org/study/summary?id=brca_metabric

https://www.ncbi.nlm.nih.gov/geo/query/acc.cgi?acc=GSE176078

https://github.com/danielyehoshua123/Centaur-phenotype-analysis

### Competing Interest Statement

The authors have declared no competing interest.

### Summary of Updates

Work revised to have a more focused story as well as to make it in-line with eLife format

